# A mathematical model of ctDNA shedding predicts tumor detection size

**DOI:** 10.1101/2020.02.12.946228

**Authors:** Stefano Avanzini, David M. Kurtz, Jacob J. Chabon, Everett J. Moding, Sharon Seiko Hori, Sanjiv Sam Gambhir, Ash A. Alizadeh, Maximilian Diehn, Johannes G. Reiter

## Abstract

Early cancer detection aims to find tumors before they progress to an incurable stage. We developed a stochastic mathematical model of tumor evolution and circulating tumor DNA (ctDNA) shedding to determine the potential and the limitations of cancer early detection tests. We inferred normalized ctDNA shedding rates from 176 early stage lung cancer subjects and calculated that a 15 mL blood sample contains on average 1.7 genome equivalents of ctDNA for lung tumors with a volume of 1 cm^3^. For annual screening, the model predicts median detection sizes between 3.8 and 6.6 cm^3^ corresponding to lead times between 310 and 450 days compared to current lung tumor sizes at diagnosis. For monthly cancer relapse testing based on 20 a priori known mutations, the model predicts a median detection size of 0.26 cm^3^ corresponding to a lead time of 150 days. This mechanistic framework can help to optimize early cancer detection approaches.

## Introduction

Early stage cancer patients are more likely to be cured than advanced stage cancer patients (*1*–*4*). For example, the five-year survival rate of lung cancer patients who are diagnosed at a localized stage is 57% while for those diagnosed with distant metastases is only 5% (*5*). Unfortunately, only 16% of lung cancers are diagnosed at a localized stage. Recently, multiple studies presented new minimally invasive approaches based on cell-free DNA (cfDNA) to detect cancer from blood samples (*6*–*15*). Most cfDNA in the bloodstream is derived from normal cells, while a small proportion is derived from tumor cells and is known as circulating tumor DNA (ctDNA). Presumably ctDNA is shed by tumor cells undergoing apoptosis or necrosis (*16*–*18*). Small tumors would therefore be harder to detect because fewer tumor cells undergo cell death.

Previous studies showed that 30-100% of symptomatic tumors (mostly larger than 3 cm^3^) can be detected from a 10 – 15 mL blood sample (*8*, *9*, *14*). However, assessing whether blood-based tests can also detect still asymptomatic tumors at sizes smaller than 3 cm^3^ with a sufficiently high specificity to reduce cancer mortality requires elaborate clinical trials with tens of thousands of participants (*19*). While such trials are already under way, we lack the mechanistic frameworks necessary to predict the expected size of tumors that would be detected with a given sequencing approach and sampling frequency. Such frameworks would enable investigators to a priori choose a sequencing and sampling strategy with the highest success probability for a given patient population. For example, how would the performance of a screening test change for a subpopulation with tumors with half as many mutations (e.g., lung cancers of non-smokers vs. smokers)? Motivated by these fundamental questions, we developed a stochastic mathematical model of cancer evolution and biomarker shedding to study the potential and the limitations of blood-based cancer early detection tests across various scenarios (*20*–*23*). This mechanistic framework will help to predict the performance of ctDNA-based tumor detection approaches and thereby inform the design of future clinical trials to find cancers earlier (*24*).

## Results

### Mathematical model of cancer evolution and ctDNA shedding

We first consider early stage lung cancers with a typical tumor volume doubling time of 181 days leading to a net growth rate of *r* = ln(2)/181 ≈ 0.4% per day (*25*). Lung cancer cells approximately divide with a birth rate of *b* = 0.14 per day (*26*), and die with a death rate of *d* = *b* − *r* = 0.136 per day (**Fig. 1A**). For now, we assume that each tumor cell releases ctDNA into the bloodstream during apoptosis with a ctDNA shedding probability of *q*_*d*_ per cell death. This assumption implies that the amount of ctDNA linearly correlates with tumor burden and the slope of the linear regression has to be 1 in logarithmic space.

**Fig. 1:**
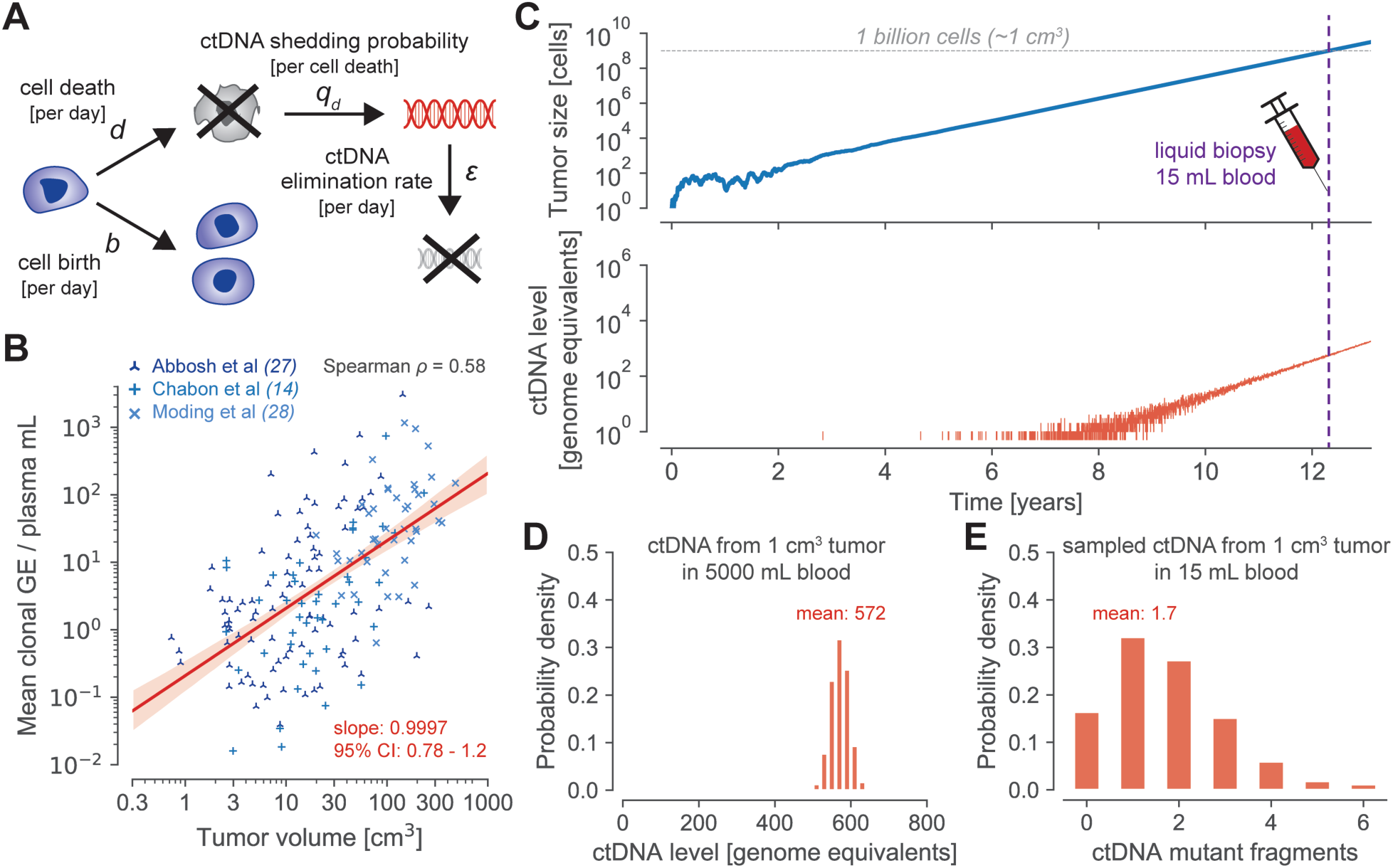
Evolutionary dynamics of ctDNA shed by a growing cancer. **A** | Mathematical model of cancer evolution and ctDNA shedding. Tumor cells divide with birth rate *b* and die with death rate *d* per day. During cell apoptosis, cells shed circulating tumor DNA (ctDNA) into the bloodstream with probability *q*_*d*_. Cell-free DNA is eliminated from the bloodstream with rate *ε* per day according to the half-life time of ctDNA, *t*_*1/2*_ = 30 minutes. **B** | Genome equivalents (GE) per plasma mL correlate with tumor volume with a slope of 0.9997 in 176 lung cancer patients (*14*, *27*, *28*). Red shaded region depicts 95% confidence interval (CI). Linear regression predicts 0.21 GE per plasma mL for a tumor volume of 1 cm^3^ leading to a shedding probability per cell death of *q*_*d*_ = 1.4 × 10^−4^ (**Methods**). **C** | A tumor starts to grow at time zero with a growth rate of *r* = *b* – *d* = 0.4% (*b* = 0.14, *d* = 0.136), typical for early-stage lung cancers (tumor doubling time of 181 days). Tumor sheds ctDNA into the bloodstream according to the product of the cell death rate *d* and the shedding probability per cell death *q*_*d*_. **D** | Distribution of ctDNA GE in the entire bloodstream at the time (purple dashed line in panel **c**) when the tumor reaches 1 billion cells (≈1 cm^3^; gray dashed line in panel **C**) leading to a mean concentration of 0.21 GE per plasma mL and a tumor fraction of 0.022% at a DNA concentration of 6.3 ng per plasma mL. **E** | Probability distribution of ctDNA mutant fragments present in a liquid biopsy of 15 mL of blood when the tumor reaches 1 billion cells (assuming the covered heterozygous mutation is present in all cancer cells).

By reanalyzing ctDNA sequencing data and tumor volumes of 176 early stage lung cancer patients of three cohorts (*14*, *27*, *28*), we found that whole genome equivalents (GE) per plasma mL indeed correlate with tumor volume with a slope of 0.9997 (red line in **Fig. 1B**; 95% confidence interval, CI: 0.78-1.2; **Methods**). We found similar linear regression slopes and intercepts in the separated three cohorts (**fig. S1**). For the combined dataset, linear regression predicted 0.21 GE per plasma mL for 1 cm^3^ of tumor volume. Based on these analyses, we inferred a mean shedding probability of *q*_*d*_ ≈ 1.4 × 10^−4^ GE per cell death (95% CI: 1.0 × 10^−4^ − 1.9 × 10^−4^ for a slope of 1). In other words, approximately 0.014% of a cancer cell’s genome is shed into the bloodstream when it undergoes apoptosis. For a ctDNA half-life time of *t*_1/2_ = 30 minutes (*16*), we calculated a ctDNA elimination rate of *ε* ≈ 33 per day. We illustrate a typical realization of this evolutionary process in **Fig. 1C** (**movie S1**). A tumor grows exponentially with a growth rate of *r* = 0.4% per day and releases ctDNA into the bloodstream with a shedding rate of *d* · *q*_*d*_. At a primary tumor size of 1 cm^3^ (∼1 billion cells (*20*)), we find on average 572 ctDNA GE circulating in the bloodstream (**Fig. 1D)**. A 15 mL blood sample contains approximately 1.7 ctDNA GE (**Fig. 1E**). At a plasma DNA concentration of 6.3 ng per plasma mL (**Methods**), 0.21 ctDNA GE per plasma mL correspond to a tumor fraction of 0.022% (assuming 6.6 pg per GE). The unit of GE can also be interpreted as the expected number of ctDNA fragments that exhibit a specific somatic heterozygous mutation. Hence, the number of ctDNA genome equivalents in a sample represents a biological limitation to detect a specific mutation in the tumor. In comparison, given a number of DNA fragments covering a specific genomic region, the number of mutated DNA fragments can be converted to a variant allele frequency (VAF) representing a technological detection limitation due to sequencing errors (*8*, *29*).

Next we aimed to calculate the expected number of ctDNA genome equivalents, *C*, circulating in the bloodstream for any tumor size *M* and derived a surprisingly simple closed-form expression. The number of ctDNA genome equivalents circulating when the tumor reaches a size of *M* cells follows a Poisson distribution with a mean of

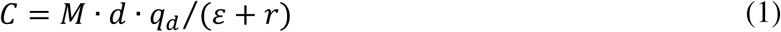

(for *M* » 1 and *d* · *q*_d_ « 1; **note S1**). For a tumor with 1 billion cells, we calculated a mean of *C* ≈ 572 ctDNA GE which perfectly matched the results from the exact computer simulations of the above defined branching process (**Fig. 1D**). Similarly, for a liquid biopsy of 15 mL blood (0.3% of 5000 mL), we calculated a mean of 1.7 GE by multiplying *C* with the fraction of the sampled blood (**Fig. 1E**).

To further demonstrate the generality of this framework and the accuracy of our analytical results, we considered tumors with different sizes, growth rates, and cell turnover rates. As expected, a tumor with 0.5 billion cells leads to roughly half the number of circulating biomarkers (*C* ≈ 286 GE; **Fig. 2A**). More surprisingly, a slowly growing lung cancer (*r* = 0.1%) leads to a significantly higher number of 585 GE than a faster growing cancer (*r* = 4%) with 420 GE at the same size of 1 cm^3^, assuming that the faster growth is achieved by a lower death rate (**Fig. 2B**). If instead the faster growth is achieved by a higher birth rate and an equal death rate, we find a smaller difference (584.2 vs. 584.9 GE). In comparison, if cancer cells divide with a birth rate of *b* = 0.25 (e.g., colorectal cancer cells (*30*); *d* = 0.246 for the same growth rate), the higher cell turnover rate would lead to an almost 2-fold (1035 vs 572 GE) increase in the amount of ctDNA in the bloodstream because of the increased rate of cells undergoing apoptosis despite the same underlying shedding probability per cell death (**Fig. 2C**). Generally, the tumor growth dynamics, the ctDNA half-life time, and the ctDNA shedding rate strongly influence ctDNA levels (**fig. S2**). The analytical results were validated by perfectly matching exact simulation results across all considered scenarios (full lines vs bars in histogram of **Fig. 2A–C**; **fig. S3**; **tables S1** and **S2**).

**Fig. 2:**
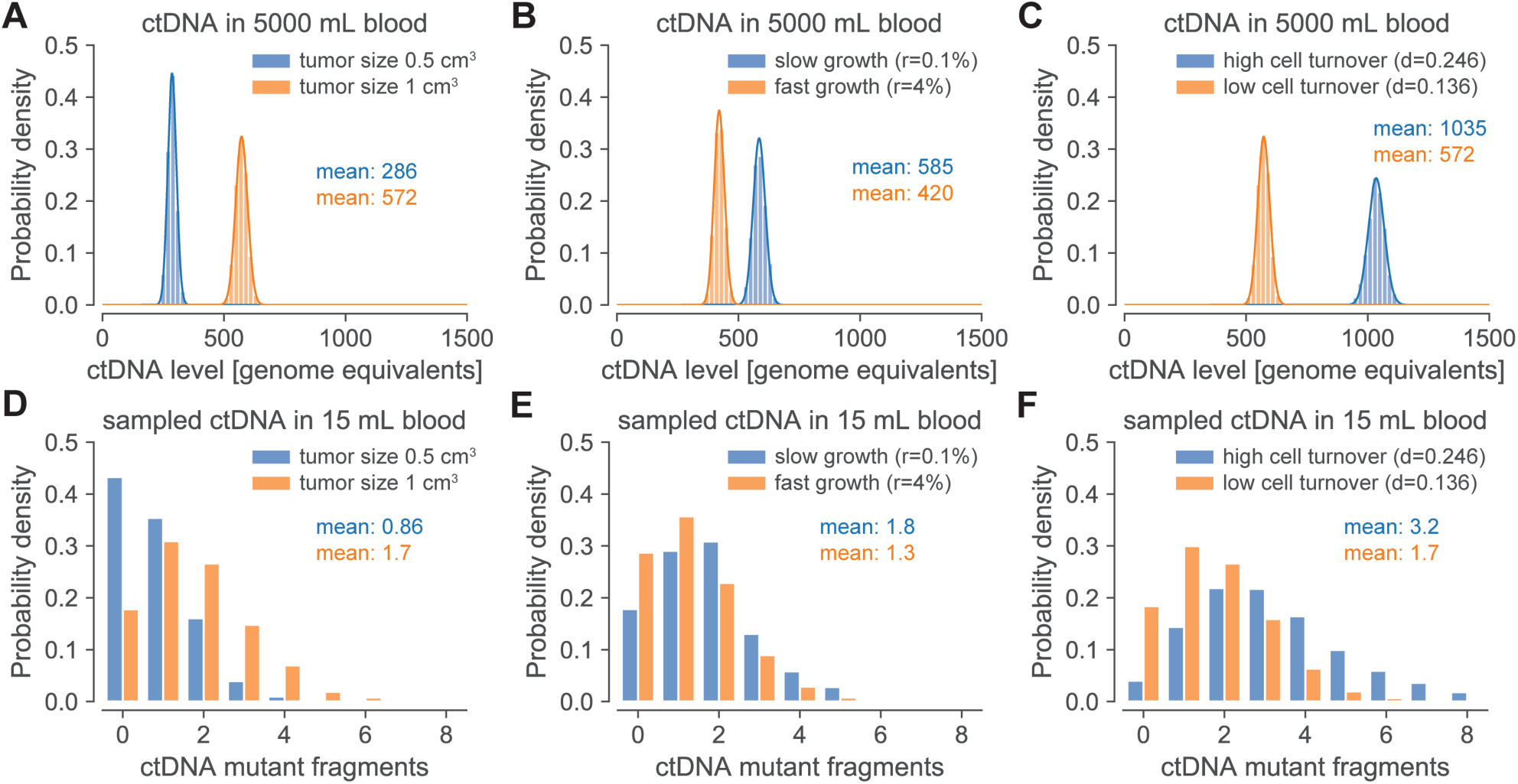
Tumor growth rate and cell turnover strongly affect the amount of ctDNA. Top panels (**A – C**) depict whole genome equivalents (GE) of ctDNA present in entire bloodstream when a lung tumor reaches a given size. Bars illustrate distribution of genome equivalents of ctDNA based on 10^4^ simulation realizations. Full lines illustrate asymptotic results (Eq. (1); **note S1**) and perfectly agree with simulation results (bars). Bottom panels (**D – F**) depict genome equivalents of ctDNA present in a 15 mL blood sample. **A** | Tumors with half the cells (0.5 vs 1 billion cells) lead to approximately half the ctDNA level in the bloodstream. **B** | Fast growing tumors lead to a lower level of ctDNA when they reach a size of 1 billion cells (≈ 1 cm^3^) because fewer cell deaths decrease the amount of released ctDNA (assuming faster growth is achieved by a lower death rate). **C** | Higher cell turnover rates lead to a higher level of ctDNA at a given tumor size (1 billion cells) compared to lower cell turnover rates because of the increased rate of cells undergoing apoptosis (if the underlying shedding probability per cell death is the same). Parameter values: birth rate *b* = 0.14 per cell per day and death rate *d* = 0.136 per cell per day (for fast growth *d* = 0.1, for slow growth *d* = 0.139); *b* = 0.25 and *d* = 0.246 for high cell turnover; ctDNA half-life time *t*_*1/2*_ = 30 minutes; ctDNA shedding probability per cell death *q*_*d*_ = 1.4 × 10^−4^ GE.

Using this mathematical framework of cancer evolution (*31*–*34*), we can predict the expected tumor detection size for an early detection test based on somatic point mutations in ctDNA (*8*, *9*). To compute realistic tumor detection sizes, we considered various sources of biological and technical errors. Given the number of wildtype and mutant fragments, we calculated the probability that a mutation arose from sequencing errors assuming a sequencing error rate of 1.5 · 10^−5^ per base-pair (*29*). To comprehend the variation of DNA concentration in plasma samples, we reanalyzed previously measured plasma DNA concentrations from Cohen et al (*9*). We used a Gamma distribution with a median of 5.2 ng of DNA per plasma mL (= 788 GE per plasma mL) to model the variability of plasma DNA concentration (**fig. S4B**; **Methods**). This analysis also revealed that plasma DNA concentrations increased in advanced cancer patients more than expected by the ctDNA amount shed from larger tumors alone (**fig. S4E**).

We distinguish two types of early cancer detection tests because of their distinct clinical use-cases and requirements: i) *cancer screening* (somatic mutations of the tumor are not known a priori) and ii) *cancer relapse detection* (somatic mutations are known a priori by sequencing a sample of the primary tumor). We start with the fundamentally simpler detection problem (ii) where the mutations are known and therefore relatively small custom sequencing panels can be used to detect a relapsing tumor.

### Tumor relapse detection if mutations are known a priori

For cancer relapse detection, we considered an aggressive lung tumor growing with *r* = 1% per day (doubling time of 69 days) and assumed a sequencing panel that covers 20 tumor-specific mutations (*27, 29, 35*–*38*). Requiring that at least one of these 20 tumor-specific mutations needs to be called as significantly present in the plasma sample to infer that the tumor relapsed, we find an AUC (area under the curve) for the ROC (receiver operating characteristic) curve of 83% for tumors with 200 mm^3^ (**Fig. 3A**). At a specificity of 99.5%, we observed a sensitivity of 15% for tumors with 200 mm^3^, assuming a sequencing efficiency of 50% (i.e. only 50% of the DNA fragments can be assessed (*14*); see **fig. S5A** for 100% sequencing efficiency). Repeatedly applying this virtual early detection test to a relapsing lung tumor led to a median detection size of 220 mm^3^ for monthly sampling and 610 mm^3^ for quarterly sampling (**fig. S5D**). Important to note is that although the same test with a specificity of 99.5% has been applied for both sampling frequencies, the monthly sampling produces 0.06 false-positive test results over 12 months of relapse testing while the quarterly sampling only produces 0.02 false-positives over 12 months. For an objective comparison, we adjusted the mutation calling thresholds such that 0.05 false-positives are expected for both sampling frequencies over 12 months of relapse testing. With this adjustment, the median detection size of quarterly testing decreased to 380 mm^3^ (**Fig. 3B**). Monthly relapse testing still led to a 32% smaller median detection size of 260 mm^3^ (see **fig. S6** for results with 100% sequencing efficiency).

**Fig. 3:**
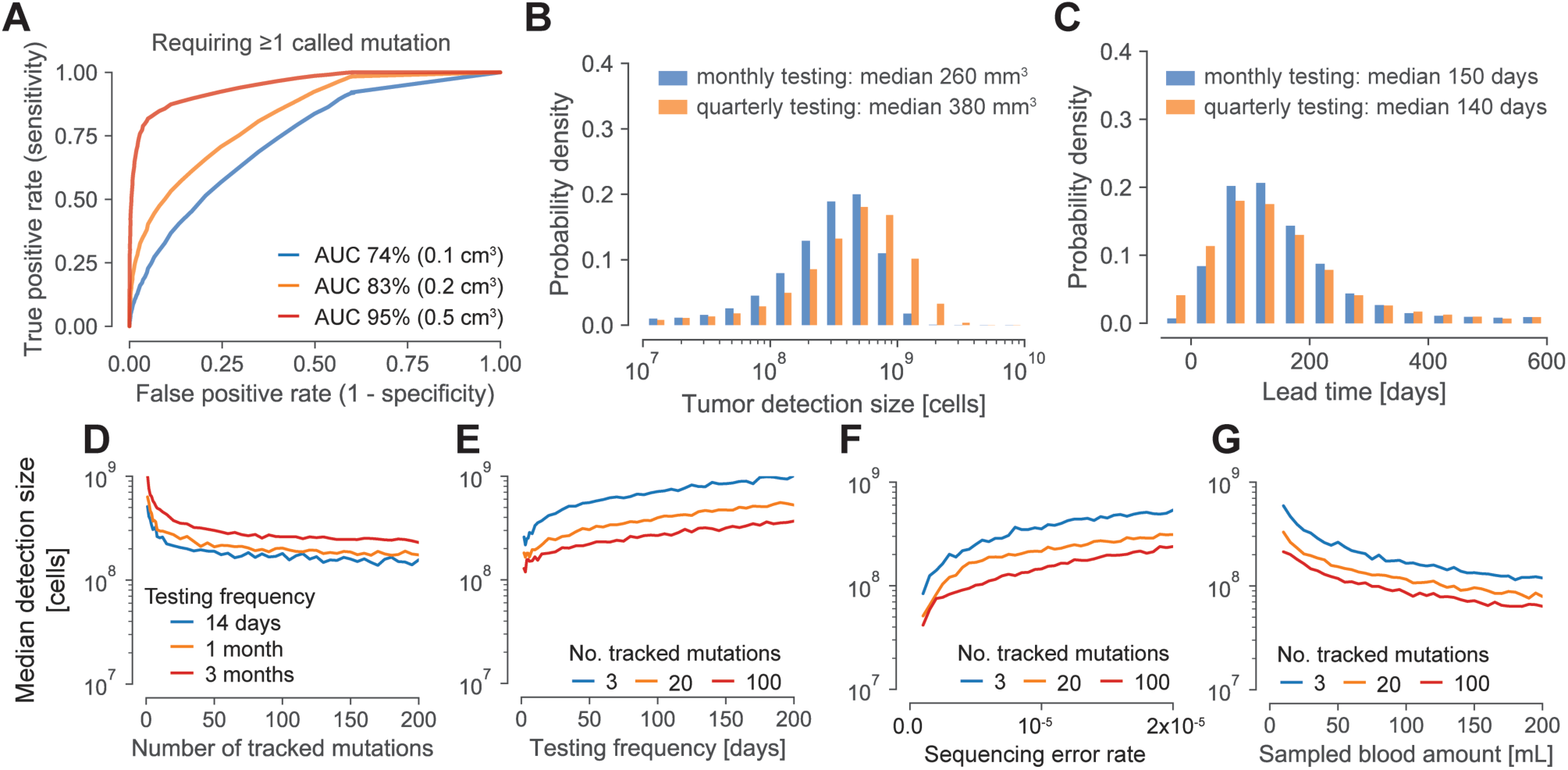
Expected tumor relapse detection size and lead time compared to current clinical relapse detection. **A** | ROC (Receiver Operating Characteristic) curves for tumors with 100 million cells (≈ 0.1 cm^3^, blue line), 200 million cells (≈ 0.2 cm^3^, orange line), or 500 million cells (≈ 0.5 cm^3^, red line) when 20 clonal tumor-specific mutations are tracked for relapse detection and one of these 20 mutations needs to be called for a positive test. AUC, area under the curve. **B – G** | For better comparability, positive detection test thresholds were set such that if the test is repeated multiple times, a combined false positive rate of 5% is obtained over all tests per year. **B** | Expected tumor detection size distributions for monthly and quarterly repeated relapse detection tests (sequencing panel covers 20 mutations). **C** | Expected lead time distributions compared to imaging-based approaches applied at the same frequency with a detection limit of 1 cm^3^ for monthly and quarterly repeated relapse detection tests (sequencing panel covers 20 clonal mutations). **D – G** | Median tumor detection sizes over the number of clonal mutations covered by the sequencing panel (panel **d**), the testing frequency (panel **E**), the sequencing error rate (panel **F**), and the sampled blood amount (panel **G**). Parameter values typical for lung cancer (if not differently specified): birth rate *b* = 0.14 per cell per day; death rate *d* = 0.13 per cell per day; ctDNA half-life time *t*_*1/2*_ = 30 minutes; ctDNA shedding probability per cell death *q*_*d*_ = 1.4 × 10^−4^ GE; sequencing efficiency of 50%; sequencing error rate per base-pair 1.5 × 10^−5^; 15 mL blood sampled per test; DNA median concentration 5.2 ng per plasma mL.

To further assess ctDNA-based relapse detection, we computed the expected lead time to imaging-based relapse detection when applied at the same frequency. We conservatively assumed a radiological detection limit of exactly 1 cm^3^ and a specificity of 100% above that limit. The median lead time of monthly ctDNA testing compared to imaging was 150 days (**Fig. 3C**). Quarterly ctDNA testing yielded a similar lead time of 140 days because of our assumption that imaging is performed at the same frequency. These predicted lead times for early stage lung cancer coincide with the reported median of ∼160 days by Chaudhuri et al. (*39*) and the slightly shorter reported median of 70 days (range, 10 – 346 days) by Abbosh et al. (*27*), likely due to two required detected mutations, less frequent relapse testing, and a lower average number of clonal mutations covered by the sequencing panel (range 3-26).

Next, we investigated how the sequencing panel size, sampling frequency, blood sample amount, sequencing error, and the number of called mutations for detection affect the tumor detection size. Although we kept the expected number of false-positives per year again constant at 0.05, the median tumor detection size strongly decreased up to a sequencing panel size of approximately 25 and then continued to minimally decrease (**Fig. 3D**). As expected, the median tumor detection size also decreased with an increasing sampling frequency (**Fig. 3E**). Weekly and more frequent relapse testing led to a large drop of the median detection size. A decreasing sequencing error rate and an increasing amount of sampled blood led to strong decreases of the expected detection sizes (**Fig. 3F**,**G**). Moreover, we found that requiring multiple called mutations for a positive detection test additionally helped to increase the sensitivity at the same specificity (*40*) (**fig. S7**).

### Tumor detection without a priori known mutations

If the mutations in the tumor are not known a priori, cancer detection becomes fundamentally more complex. Two major considerations for ctDNA-based cancer early detection are the expected number of mutations per tumor covered by the sequencing panel and the underlying sequencing error rate per base-pair of the assay. The expected number of somatic mutations covered by the sequencing panel can be maximized by focusing on recurrently mutated regions of the genome such that many more mutations per sequenced Mega-base are observed than expected from the average lung cancer mutation frequency of ∼10 mutations per Mega-base. For example, CAPP-Seq (spanning ∼300,000 base-pairs) and CancerSEEK (spanning ∼4,500 base-pairs) cover on average 9.1 and 1.1 mutations per early stage lung cancer in the TRACERx cohort (*9*, *27*, *29*) (**table S3**).

We again consider an early stage lung tumor growing with *r* = 0.4% per day. For a sequencing panel covering on average one mutation per lung cancer across 4,500 base-pairs, we computed sensitivities of 4.3%, 17%, and 54% at a specificity of 99% for tumors at sizes of 1 cm^3^, 2 cm^3^, and 4 cm^3^, respectively. Repeating this virtual early detection test annually for a growing tumor, we obtained a median detection size of 6.6 cm^3^ (diameter of 2.3 cm; **Fig. 4A**). This detection size would be 71% smaller than the current median detection size of approximately 22.5 cm^3^ (diameter of 3.5 cm; assuming that tumors are approximately spherical) for lung cancers reported in the SEER database from 2005 to 2015 (*41*). Comparing the computed detection size and the SEER median size at diagnosis, we calculated a lead time to current diagnosis times of 310 days for a typical growth rate of early stage lung cancer (**Fig. 4D**). For 4.4% of cancers, we observed a negative lead time. In other words, those cancers became symptomatic (i.e., reached typical diagnosis size of 22.5 cm^3^) before they were detected by screening. Faster growing tumors led to larger detection sizes. For example, for more than twice as fast-growing tumors (*r* = 1%, tumor volume doubling time of 69 days), we computed a median detection size by screening of 21 cm^3^ for annual screening (**fig. S8A**). In this case, only 52% of cancers were detected by screening and 48% of cancers became symptomatic before being detected by screening leading to a median screening detection size of 8.1 cm^3^. Note that these tumors grow from 0.1 cm^3^ to 3.8 cm^3^ in just one year and are therefore very hard to detect with a screening program.

**Fig. 4:**
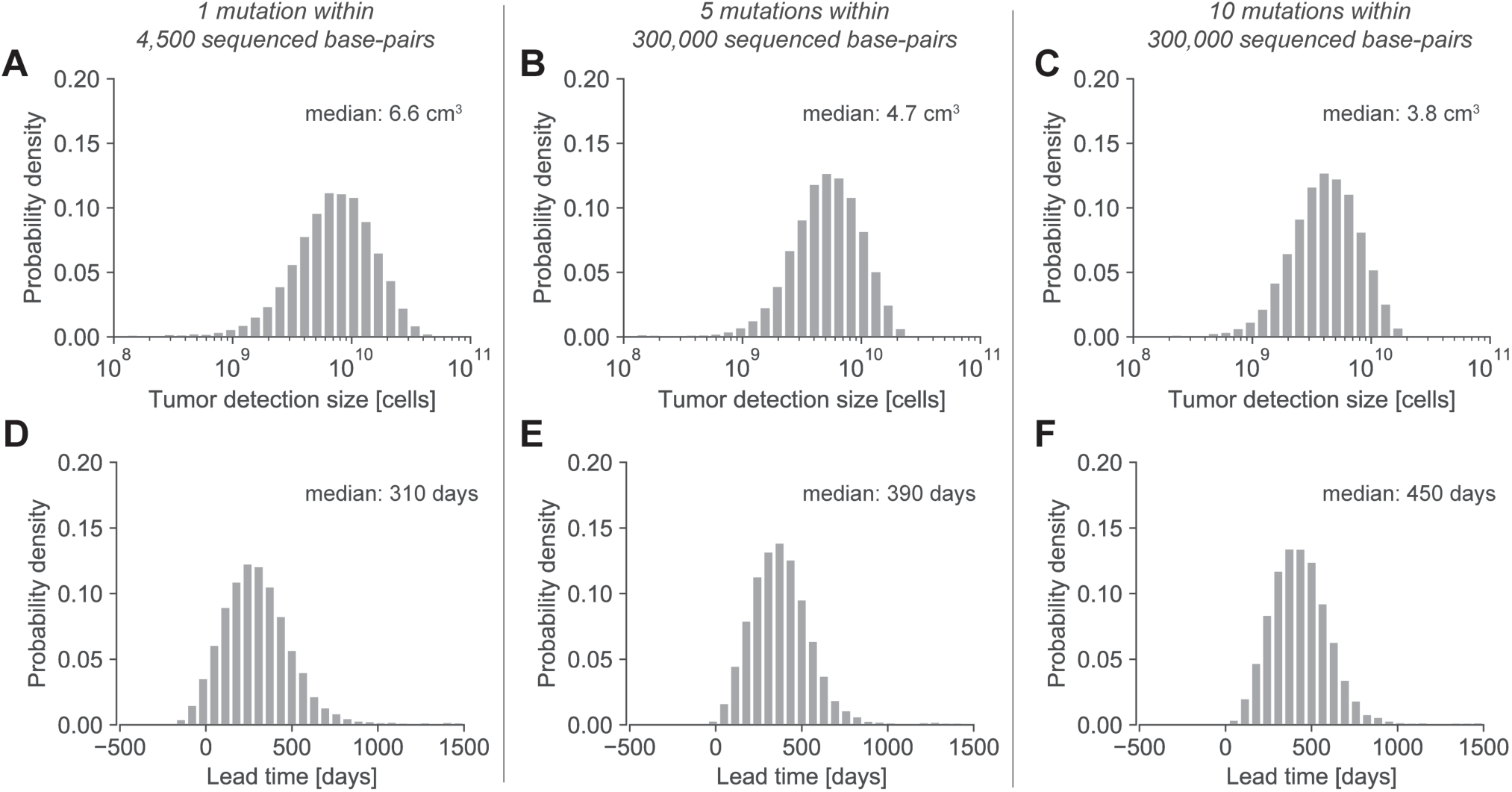
Expected tumor detection size and lead time distributions for different screening scenarios. **A – C** | Tumor detection size distributions for annually repeated virtual screening tests for different sequencing panels and number of covered mutations for cancers of never-smoking subjects (panel **B**) and subjects with a smoking history (panel **C**). Number of expected mutations covered by the smaller panel is similar for cancers of smokers and non-smokers (panel **A**; **table S3**). **D – F** | Lead time distributions for annually repeated virtual screening tests compared to current clinical diagnosis times calculated from detection sizes in the SEER database, assuming an early stage lung cancer growth rate of 0.4% per day. Parameter values typical for early stage lung cancer: 99% test specificity; birth rate *b* = 0.14 per cell per day; death rate *d* = 0.136 per cell per day; ctDNA half-life time *t*_*1/2*_ = 30 minutes; ctDNA shedding probability per cell death *q*_*d*_ = 1.4 × 10^−4^; sequencing error rate per base-pair 1.5 × 10^−5^; sequencing efficiency 50%; 15 mL blood sampled per test; DNA median concentration 5.2 ng per plasma mL.

In comparison, for a sequencing panel covering 300,000 base-pairs (e.g., CAPP-Seq), we computed sensitivities of 29% and 47% at a specificity of 99% for tumors with a size of 2 cm^3^ in never-smoking subjects and in subjects with a smoking history, respectively. We separately analyzed smokers and non-smokers because in contrast to the above evaluated panel which focused on driver gene mutations, the expected number of mutations covered by the larger panel for cancers of never-smoking subjects was lower than for cancers of subjects with a smoking history (5 vs. 10 mutations; **table S3**). For an annually repeated screening test, the expected median detection size was 4.7 cm^3^ (diameter of 2.1 cm) for lung tumors of never-smokers and 3.8 cm^3^ (diameter of 1.9 cm) for tumors of subjects with a smoking-history (**Fig. 4B**,**C**). These detection sizes correspond to lead times compared to current tumor sizes at diagnosis of 390 and 450 days for tumors of never-smokers and smokers, respectively (**Fig. 4E**,**F**). Similarly high lead times of up to 450 days were observed in the CCGA (Circulating Cell-free Genome Atlas) study (*15*).

For the above calculations, we assumed that the detected mutations in cfDNA were unique to cancers. However, expanded clones in blood or in normal tissue and benign lesions frequently exhibit somatic mutations (*42*–*44*), which can be observed in the cfDNA (*14*, *45*) and hence hamper the specificity of a cancer screening test. We therefore assumed that white blood cells are sequenced separately to remove somatic mutations arising due to clonal hematopoiesis. Moreover, previous studies showed that on average eight benign lung nodules with a diameter of ∼4 mm exist in subjects undergoing lung cancer screening (*46*). Because most benign lesions are smaller than malignant tumors and since benign cells typically replicate with a lower rate than malignant cells, they are expected to release comparatively less ctDNA into the bloodstream (*26*). Our calculations suggest that these nodules would only slightly increase the expected detection size – even if benign cells exhibit the same ctDNA shedding probability and somatic mutation load as malignant cells (**fig. S9**).

While these predicted tumor detection sizes are encouraging, a major challenge for every cancer detection test is the low incidence rate of cancers. Relatively common cancers such as lung cancer have yearly incidence rates of approximately 1 in 2,000 (*5*). Because more than 98% of lung cancers occur in people above 50 (∼30% of the US population), the incidence rate increases to 1 in 600 in this age group. Hence, approximately six false-positives would be expected for one true-positive at a test specificity of 99%. We calculated a PPV (positive predictive value) of 2.7% and an NPV (negative predictive value) of 99.9% for a lung tumor with 2 billion cells using a sequencing panel of 4,500 base-pairs as in **Fig. 4A**. For heavy smokers in their late sixties the incidence rate increases to 1 in 120 and the same test would have a PPV of 12% and an NPV of 99.3% for lung cancers with 2 billion cells. In comparison, a PPV of 3.8% was reported for low-dose CT lung cancer screening (*47*).

### ctDNA shedding during apoptosis, necrosis, and proliferation

So far we assumed that ctDNA shedding occurs exclusively during apoptosis. We can generalize our framework such that the effective shedding rate *λ* is given by the sum of ctDNA fragments shed during apoptosis, necrosis, and proliferation: *λ* = *d* · *q_d_* + *q_n_* + *b* · *q_b_* where *q_b_* denotes the shedding probability per cell division and *q_n_* denotes the shedding rate from necrosis per unit of time (*10*, *16*). We show that independent of the three shedding processes, the amount of ctDNA when the tumor reaches a size of *M* cells remains approximately Poisson-distributed with a mean of *C* = *M* · *λ*/(*ε* + *r*) (assuming *M* » 1 and *λ* « 1; **note S1**) where the effective shedding rate represents the sum of the operating ctDNA shedding processes (**fig. S10**; **table S1**).

## Discussion

Our mathematical framework provides a theoretical upper-bound for the performance of mutation-based ctDNA cancer early detection tests. A critical parameter for these estimates is the normalized ctDNA shedding rate per cancer cell. Based on ctDNA data of early stage lung cancer (*14*, *27*, *28*), the stochastic model suggests that decreasing the sequencing error rate, increasing the amount of sampled plasma, increasing the sequencing panel size, and increasing the sampling rate can drastically decrease the expected tumor relapse detection size at the same normalized annual false-positive rate (**Fig. 3**). The virtual screening computations indicate that lung tumors would be detected when they reach a diameter of 2.3 cm in an annual screening program with a sequencing panel of 4,500 base-pairs (**Fig. 4A**). According to the lung cancer staging system for tumor sizes, 11% of the detected cases would be classified as T1a (≤1 cm), 21% as T1b (>1 but ≤2 cm), and 52% as T1c (>2 but ≤3 cm) (*48*). Only 16% of the detected tumors would have reached sizes for stage T2a (>3 but ≤4 cm) or beyond. Although these calculations suggest that most tumors can only be detected when they reach sizes of billions of cells, detecting some tumors before they become symptomatic and shifting some diagnosis to an earlier stage can have an enormous impact on cancer mortality (*4*, *49*).

This study has several limitations. First, our understanding of ctDNA shedding and its variance across lung tumors and other tumor types remains limited. More studies such as those by Abbosh et al, Chabon et al, and Moding et al correlating ctDNA levels with tumor volume and other clinicopathological features in lung cancer patients are required to inform shedding rate inferences in other tumor types (*14*, *27*, *28*). The high concordance both in the slope and the intercept of the linear regression analysis across the three aforementioned lung cancer cohorts is reassuring (**Fig. 1B**; **fig. S1**). Other biological factors such as tumor histology and stage additionally influence ctDNA levels (*14*) and could be included in future models when more data becomes available. Second, the ctDNA shedding dynamics of precursor lesions and how their presence interferes with cancer early detection are largely unknown (**fig. S9**). Third, our analysis was limited to point mutations present in ctDNA. Including additional cancer-associated characteristics of ctDNA or other biomarkers can help to further decrease the expected detection size (*9*–*15, 17*). Last, we verified our mathematical results through exact computer simulations, however, the predicted tumor detection sizes will need to be validated in large clinical studies. According to the model, every tumor would eventually become detectable – but some will become symptomatic first (e.g., those cancers with negative lead times in **Fig. 4**). We note that tumor-specific ctDNA mutant fragments were detected in all TRACERx subjects but no mutations were called due to the high specificity requirements in 40% (38/96) of subjects (*27*).

A major challenge of cancer early detection is the stochastic nature of cancer initiation and progression. While new technologies can detect smaller and smaller tumors, the optimal treatment of asymptomatic tumors is often unclear and needs to be balanced with the risk for overtreatment as well as undertreatment (*3*, *4*). For example, in more than 20% of smokers suspicious lesions can be found by low-dose computed tomographic screening, nevertheless, lung cancer was detected only in ∼0.6% of screened smokers within a 10-year follow-up (*50*, *51*). Similarly, around 33% of humans harbor precursor lesions in their pancreas but the life-time risk of developing pancreatic cancer is 1.6% (*41*, *52*). Hence, most precursor lesions do not progress to cancer within the life-time of humans (*44*, *53*). Because cancer incidence strongly depends on factors such as age, genetic predisposition, life-style, or exposure to mutagens (e.g., sun, smoke, etc.), screening programs often focus on high-risk individuals to decrease the chances of overtreatment. Our results show that cancer screening and surveillance strategies can be further optimized and personalized by comprehensive mathematical models of cancer evolution and biomarker shedding (*21, 53*–*58*).

## Methods

### Parameter selection and inference

We used previously measured values of cell division rates and tumor volume doubling times to obtain the tumor growth rates and death rates. The estimated average time between cell divisions of lung cancer cells is 7 days (*26*), resulting in a cell birth rate of *b* = 0.14 per day. Given a volume doubling time of ∼180 days of stage I lung cancers (*25*), we find a tumor growth rate *r* = ln(2)/180 ≈ 0.4% per day and a death rate of *d* = *b* − *r* = 0.136 per day. For the analysis of tumors with a high cell turnover rate (**Fig. 2C**), we assumed an average time between cell divisions of colorectal cancer cells of 4 days (*26*), resulting in a cell birth rate of *b* = 0.25 per day. For the analysis with co-existing benign lesions (**fig. S9**), we explored the effects of nodules at fixed sizes due to their doubling times of ≥ 500 days and assumed that cells in benign lung nodules replicate with rates of *b*_bn_ = *d*_bn_ = 0.07 per day (*25*, *26*).

To estimate the ctDNA shedding rate per cancer cell, we reanalyzed data from 176 early-stage lung cancer patients studied in Chabon et al (*14*), Abbosh et al (*27*), and Moding et al (*28*). In Chabon et al, metabolic tumor volumes were computed from whole body ^18^F-FDG PET (positron emission tomography)-CT scans in 81 subjects. For 46 of these subjects, tumor-specific ctDNA mutations were reported. Reanalyzed gross tumor volume from Abbosh et al and Moding et al were estimated from preoperative CT scans. Genome equivalents (GE) per plasma mL were calculated from the mean variant allele frequency (VAF) times the plasma cfDNA concentration in a subject divided by 0.0033 ng (weight of haploid human genome). Estimates can be found in Supplementary Table 3 of Chabon et al and Supplementary Table 2 of Moding et al. For the data in Abbosh et al, the mean VAF of somatic mutations was calculated across all mutations (including those that were not called in the liquid biopsy) that were identified as clonal in the primary tumor (in subject CRUK0053, all mutations were considered because no clonality information was available; see Supplementary Table 5 of ref. (*27*)).

To infer the shedding rate *λ*, we used the predicted 0.21 GE per plasma mL for a tumor with a volume of 1 cm^3^ (**Fig. 1B**) and set up the following equilibrium equation of *λ* · 10^9^ cells per volume mL = 577.5 · *ε*, assuming 577.5 (= 0.21 · 5000 · 55%) whole genome equivalents in 5000 mL of blood with a plasma concentration of 55%. We followed previous estimates of the ctDNA half-life time of *t*_1/2_ = 30 minutes and explored the effects of half-life times from 20 to 120 minutes (*16*) (**fig. S2**). We calculated the ctDNA elimination rate as 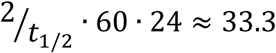 fragments per day. We found a shedding rate of *λ* = 1.9 · 10^−5^ GE per cell per day leading to a shedding probability of 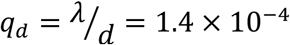 GE per cell death, assuming that ctDNA is exclusively shed during cell apoptosis. To estimate the shedding rate of cells in other tissue types, we can rescale the shedding probability by the corresponding cell death rate. For example, we estimate the shedding rate of cells in benign lung nodules as *λ*_bn_ = *d*_bn_ · *q*_d_ ≈ 9.8 · 10^−6^ GE per cell per day.

### In silico sampling and sequencing of plasma DNA

To compare different screening strategies, we employed virtual sampling and sequencing of DNA. For each liquid biopsy, ctDNA fragments in 15 mL of blood were sampled from 5000 mL of blood according to a binomial distribution. For the virtual detection tests in **Figs. 3, 4** and **figs. S5-S9**, plasma cfDNA concentrations were sampled from a Gamma distribution (mean and median of 6.3 and 5.2 ng per plasma mL) as illustrated in **fig. S4B**. We assumed that 50% of cfDNA fragments are assessed by sequencing (except in **figs. S5A-C** and **S6**). To calculate the cfDNA GE, we assumed a weight of 6.6 pg per GE (*17*). The sequencing error rate per base-pair was set to *e*_seq_ = 1.5 · 10^−5^ (*8*, *29*). For each mutation covered by the sequencing panel, we account for sequencing errors by calculating a *P* value as 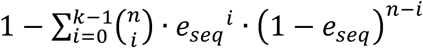 where *k* denotes the number of observed fragments supporting the mutation and *n* denotes the total number of fragments covering the mutation’s genomic position.

## Acknowledgments

We thank Tibor Antal, David Cheek, and Michael Nicholson for valuable discussions and feedback. Some of the computing for this project was performed on the Stanford Sherlock cluster. We would like to thank Stanford University and the Stanford Research Computing Center for providing computational resources and support that contributed to these research results.

## Author contributions

J.G.R. conceived and supervised the study. S.A. and J.G.R. developed the mathematical model and performed the computational analysis. S.A., D.M.K., J.J.C., E.J.M., A.A.A., M.D., J.G.R. analyzed data. All authors discussed and interpreted data. J.G.R. and S.A. wrote the original manuscript. All authors reviewed and edited the manuscript.

## References

1. R. Etzioni et al., Early detection: The case for early detection. Nat. Rev. Cancer. 3, 243 (2003).

2. M. Song, B. Vogelstein, E. L. Giovannucci, W. C. Willett, C. Tomasetti, Cancer prevention: Molecular and epidemiologic consensus. Science. 361, 1317–1318 (2018).

3. S. Srivastava et al., Cancer overdiagnosis: a biological challenge and clinical dilemma. Nat. Rev. Cancer. 19, 349–358 (2019).

4. A. K. Mattox et al., Applications of liquid biopsies for cancer. Sci. Transl. Med. 11, eaay1984 (2019).

5. R. L. Siegel, K. D. Miller, A. Jemal, Cancer Statistics, 2020. CA. Cancer J. Clin. 70, 7–30 (2020).

6. C. Bettegowda et al., Detection of Circulating Tumor DNA in Early- and Late-Stage Human Malignancies. Sci. Transl. Med. 6, 224ra24 (2014).

7. A. M. Newman et al., An ultrasensitive method for quantitating circulating tumor DNA with broad patient coverage. Nat. Med. 20, 548 (2014).

8. J. Phallen et al., Direct detection of early-stage cancers using circulating tumor DNA. Sci Transl Med. 9, eaan2415 (2017).

9. J. D. Cohen et al., Detection and localization of surgically resectable cancers with a multi-analyte blood test. Science. 357, 926–930 (2018).

10. F. Mouliere et al., Enhanced detection of circulating tumor DNA by fragment size analysis. Sci. Transl. Med. 10, eaat4921 (2018).

11. S. Y. Shen et al., Sensitive tumour detection and classification using plasma cell-free DNA methylomes. Nature. 563, 579 (2018).

12. S. Cristiano et al., Genome-wide cell-free DNA fragmentation in patients with cancer. Nature. 570, 385–389 (2019).

13. P. Ulz et al., Inferring expressed genes by whole-genome sequencing of plasma DNA. Nat. Genet. 48, 1273 (2016).

14. J. J. Chabon et al., Integrating genomic features for non-invasive early lung cancer detection. Nature (2020), doi: 10.1038/s41586-020-2140-0.

15. M. C. Liu et al., Sensitive and specific multi-cancer detection and localization using methylation signatures in cell-free DNA. Ann. Oncol. (2020), doi: 10.1016/j.annonc.2020.02.011.

16. J. C. M. Wan et al., Liquid biopsies come of age: towards implementation of circulating tumour DNA. Nat. Rev. Cancer. 17, 223–238 (2017).

17. E. Heitzer, I. S. Haque, C. E. S. Roberts, M. R. Speicher, Current and future perspectives of liquid biopsies in genomics-driven oncology. Nat. Rev. Genet. 20, 71–88 (2019).

18. H. Schwarzenbach, D. S. B. Hoon, K. Pantel, Cell-free nucleic acids as biomarkers in cancer patients. Nat. Rev. Cancer. 11, 426–437 (2011).

19. A. M. Aravanis, M. Lee, R. D. Klausner, Next-generation sequencing of circulating tumor DNA for early cancer detection. Cell. 168, 571–574 (2017).

20. S. S. Hori, S. S. Gambhir, Mathematical model identifies blood biomarker--based early cancer detection strategies and limitations. Sci. Transl. Med. 3, 109ra116 (2011).

21. S. S. Hori, A. M. Lutz, R. Paulmurugan, S. S. Gambhir, A Model-Based Personalized Cancer Screening Strategy for Detecting Early-Stage Tumors Using Blood-Borne Biomarkers. Cancer Res. 77, 2570–2584 (2017).

22. B. Vogelstein et al., Cancer Genome Landscapes. Science. 339, 1546–1558 (2013).

23. M. Greaves, C. C. Maley, Clonal evolution in cancer. Nature. 481, 306–313 (2012).

24. D. W. Cescon, S. V Bratman, S. M. Chan, L. L. Siu, Circulating tumor DNA and liquid biopsy in oncology. Nat. Cancer. 1, 276–290 (2020).

25. H. T. Winer-Muram et al., Volumetric growth rate of stage I lung cancer prior to treatment: serial CT scanning. Radiology. 223, 798–805 (2002).

26. D. A. Rew, G. D. Wilson, Cell production rates in human tissues and tumours and their significance. Part II: clinical data. Eur. J. Surg. Oncol. 26, 405–417 (2000).

27. C. Abbosh et al., Phylogenetic ctDNA analysis depicts early-stage lung cancer evolution. Nature. 545, 446–451 (2017).

28. E. J. Moding et al., Circulating tumor DNA dynamics predict benefit from consolidation immunotherapy in locally advanced non-small-cell lung cancer. Nat. Cancer. 1, 176–183 (2020).

29. A. M. Newman et al., Integrated digital error suppression for improved detection of circulating tumor DNA. Nat. Biotechnol. 34, 547 (2016).

30. I. Bozic et al., Accumulation of driver and passenger mutations during tumor progression. Proc Natl Acad Sci USA. 107, 18545–18550 (2010).

31. R. Durrett, Branching process models of cancer (Springer, 2015).

32. N. Beerenwinkel, R. F. Schwarz, M. Gerstung, F. Markowetz, Cancer evolution: mathematical models and computational inference. Syst. Biol. 64, e1–e25 (2015).

33. P. M. Altrock, L. L. Liu, F. Michor, The mathematics of cancer: integrating quantitative models. Nat. Rev. Cancer. 15, 730–745 (2015).

34. D. Wodarz, N. L. Komarova, Dynamics of Cancer: Mathematical Foundations of Oncology (World Scientific Publishing, Singapore, 2014).

35. B. R. McDonald et al., Personalized circulating tumor DNA analysis to detect residual disease after neoadjuvant therapy in breast cancer. Sci. Transl. Med. 11, eaax7392 (2019).

36. J. Tie et al., Circulating tumor DNA analysis detects minimal residual disease and predicts recurrence in patients with stage II colon cancer. Sci. Transl. Med. 8, 346ra92 (2016).

37. T. Reinert et al., Analysis of plasma cell-free DNA by ultradeep sequencing in patients with stages I to III colorectal cancer. JAMA Oncol. (2019), doi: 10.1001/jamaoncol.2019.0528.

38. K. H. Khan et al., Longitudinal liquid biopsy and mathematical modeling of clonal evolution forecast time to treatment failure in the PROSPECT-C phase II colorectal cancer clinical trial. Cancer Discov. 8, 1270–1285 (2018).

39. A. A. Chaudhuri et al., Early detection of molecular residual disease in localized lung cancer by circulating tumor DNA profiling. Cancer Discov. 7, 1394–1403 (2017).

40. J. C. M. Wan et al., High-sensitivity monitoring of ctDNA by patient-specific sequencing panels and integration of variant reads. bioRxiv, 759399 (2019).

41. NCI Seer, Surveillance, Epidemiology, and End Results (SEER) Research Data 1973-2015 - ASCII Text Data (2018), (available at https://seer.cancer.gov).

42. I. Martincorena, P. J. Campbell, Somatic mutation in cancer and normal cells. Science 349, 1483–1489 (2015).

43. A. Yokoyama et al., Age-related remodelling of oesophageal epithelia by mutated cancer drivers. Nature. 565, 312–317 (2019).

44. A. P. Makohon-Moore et al., Precancerous neoplastic cells can move through the pancreatic ductal system. Nature. 561, 201–205 (2018).

45. P. Razavi et al., High-intensity sequencing reveals the sources of plasma circulating cell-free DNA variants. Nat. Med. 25, 1928–1937 (2019).

46. A. McWilliams et al., Probability of cancer in pulmonary nodules detected on first screening CT. N. Engl. J. Med. 369, 910–919 (2013).

47. The National Lung Screening Trial Research Team, Results of Initial Low-Dose Computed Tomographic Screening for Lung Cancer. N. Engl. J. Med. 368, 1980–1991 (2013).

48. F. C. Detterbeck, D. J. Boffa, A. W. Kim, L. T. Tanoue, The eighth edition lung cancer stage classification. Chest. 151, 193–203 (2017).

49. C. A. Clarke et al., Projected Reductions in Absolute Cancer{\textendash}Related Deaths from Diagnosing Cancers Before Metastasis, 2006{\textendash}2015. Cancer Epidemiol. Prev. Biomarkers (2020), doi: 10.1158/1055-9965.EPI-19-1366.

50. The National Lung Screening Trial Research Team, Reduced lung-cancer mortality with low-dose computed tomographic screening. N. Engl. J. Med. 365, 395–409 (2011).

51. H. J. de Koning et al., Reduced Lung-Cancer Mortality with Volume CT Screening in a Randomized Trial. N. Engl. J. Med. (2020), doi: 10.1056/NEJMoa1911793.

52. A. P. Makohon-Moore, C. A. Iacobuzio-Donahue, Pancreatic cancer biology and genetics from an evolutionary perspective. Nat. Rev. Cancer. 16, 553–565 (2016).

53. B. M. Lang, J. Kuipers, B. Misselwitz, N. Beerenwinkel, Predicting colorectal cancer risk from adenoma detection via a two-type branching process model. PLOS Comput. Biol. 16, e1007552 (2020).

54. M. E. Hochberg, F. Thomas, E. Assenat, U. Hibner, Preventive evolutionary medicine of cancers. Evol. Appl. 6, 134–143 (2013).

55. H. J. de Koning et al., Benefits and harms of computed tomography lung cancer screening strategies: a comparative modeling study for the US Preventive Services Task Force. Ann. Intern. Med. 160, 311–320 (2014).

56. K. Curtius, A. Dewanji, W. D. Hazelton, J. H. Rubenstein, E. G. Luebeck, Optimal timing for cancer screening and adaptive surveillance using mathematical modeling. bioRxiv (2020), doi: 10.1101/2020.02.11.927475.

57. M. D. Ryser et al., Outcomes of active surveillance for ductal carcinoma in situ: a computational risk analysis. J. Natl. Cancer Inst. 108, djv372 (2016).

58. K. Lahouel et al., Revisiting the tumorigenesis timeline with a data-driven generative model. Proc Natl Acad Sci USA. 117, 857–864 (2020).

