## Supplementary Information for "A mathematical model of ctDNA shedding predicts tumor detection size"

#### Contents

**Supplementary Figures** **2**

**Supplementary Tables** **12**

**Supplementary Note S1: Mathematical Modeling** **15**

**References** **41**

### Supplementary Figures

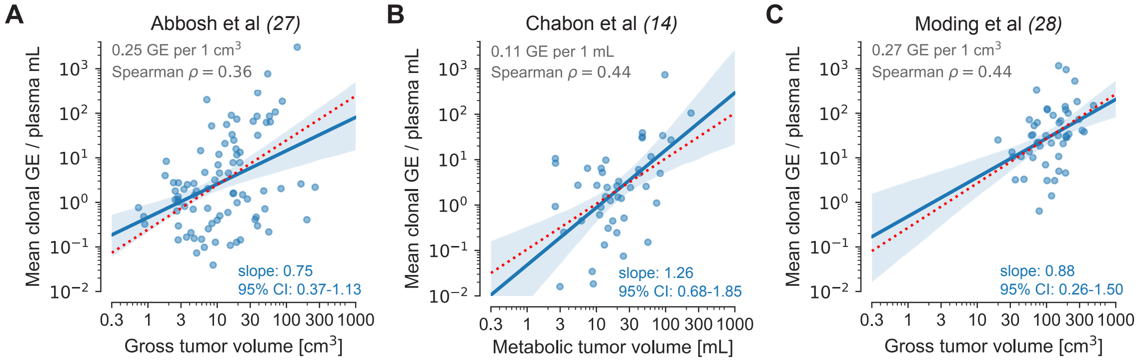


**Fig. S1: Amount of ctDNA correlates with tumor volume in early stage lung cancer.** Genome equivalents (GE) correlate with tumor volume with a linear regression slope of close to 1 (red dotted line) in logarithmic space. Gross tumor volumes were inferred from CT-scans. Shaded region depicts the 95% confidence interval of a linear regression in log-log space. **A** | Reanalyzed data of 87 early stage non-small-cell lung cancer patients of the TRACERx cohort where accurate volumetric estimates were availalbe (*27*). Genome equivalents were calculated from the mean variant allele frequency (VAF) of all clonal mutations per subject (see **Methods** for details). For subject CRUK0053, clonality estimates were not available and hence all mutations were assumed to be clonal. If tumors with lower volumes (close to the limit of detection with higher variance) are excluded, the slope approaches 1. Excluding tumors with volumes below 2 and 3 cm$^3$ leads to slopes of 0.85 and 0.91, respectively. Linear regression with a fixed slope of 1 predicts 0.25 GE per plasma mL for 1 cm$^3$ of tumor volume (95% CI: $0.15 - 0.40$ GE per plasma mL). **B** | Reanalyzed data of 46 early stage non-small-cell lung cancer patients (*14*). Genome equivalents were reported in Supplementary Table 3 of the original study. Linear regression with a fixed slope of 1 predicts 0.11 GE per plasma mL for 1 mL of tumor volume (95% CI: $0.055 - 0.21$ GE per plasma mL). **C** | Reanalyzed data of 49 locally advanced non-small-cell lung cancer patients (*28*). Genome equivalents were reported in Table S2 of the original study. Note that only for 6 subjects, mutations were known a priori. For the remaining 43 subjects, mutation calls were based on the blood samples and hence required stricter calling thresholds decreasing the sensitivity. Linear regression with a fixed slope of 1 predicts 0.27 GE per plasma mL for 1 mL of tumor volume (95% CI: $0.18 - 0.41$ GE per plasma mL). Six subjects were both part of the cohorts of refs. (*14*) and (*28*) and only kept once in the combined dataset.

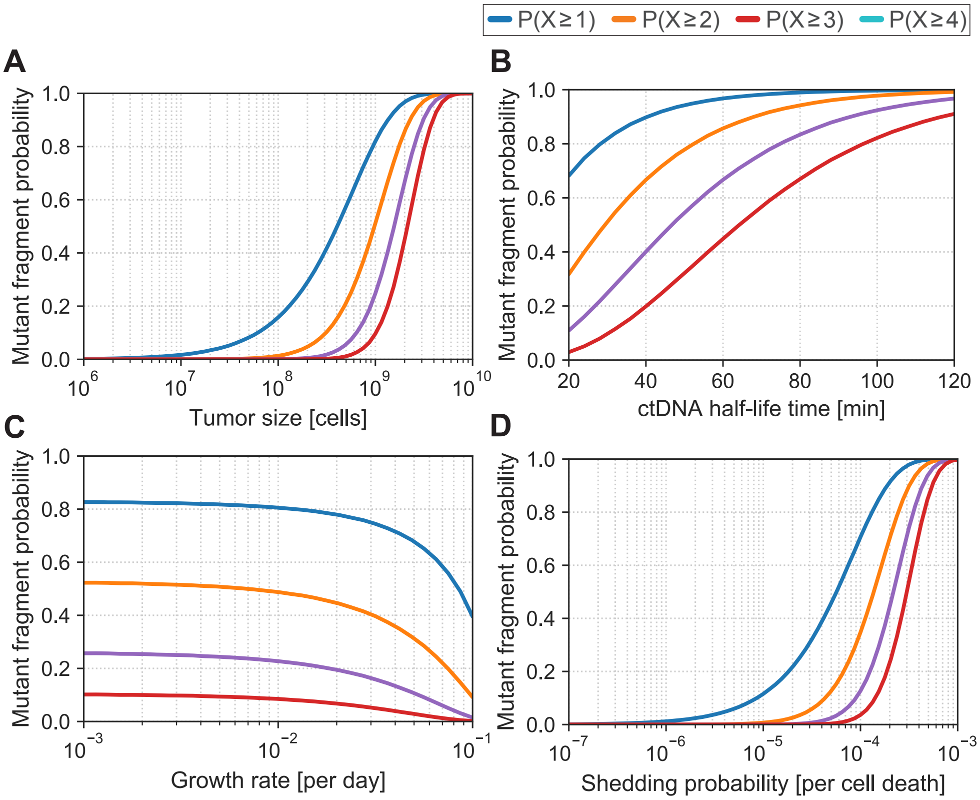


**Fig. S2: Detection probability of ctDNA fragments in a 15 mL liquid biopsy at given tumor size depends on many parameters.** Full lines denote the probability to find at least 1 (blue), 2 (orange), 3 (red), or 4 (turquoise) mutant ctDNA fragments in a liquid biopsy of 15 mL blood. Probability of finding mutant ctDNA fragments of a specific region corresponds to diploid genome equivalents. Standard parameter values (if not otherwise noted): birth rate $b = 0.14$ per cell per day, death rate $d = 0.136$ per cell per day, tumor detection size $M = 10^9$, ctDNA half-life time $t_{1/2} = 30$ minutes, ctDNA shedding probability per cell death $q_d = 1.4 \cdot 10^{-4}$ (**Methods**). **A** | Tumors with fewer than $10^9$ cells ($\approx 1$ cm$^3$) rarely shed sufficient ctDNA that individual somatic mutations can be robustly detected. **B** | Higher ctDNA half-life time $t_{1/2}$ increases the number of mutant ctDNA fragments at a given tumor size. **C** | Slower growing tumors lead to higher numbers of mutant ctDNA fragments compared with fast growing tumor at the same size. Growth rate is varied by changing the death rate and keeping the birth rate constant. **D** | A slightly increased ctDNA shedding probability $q_d$ boosts the detection probability.

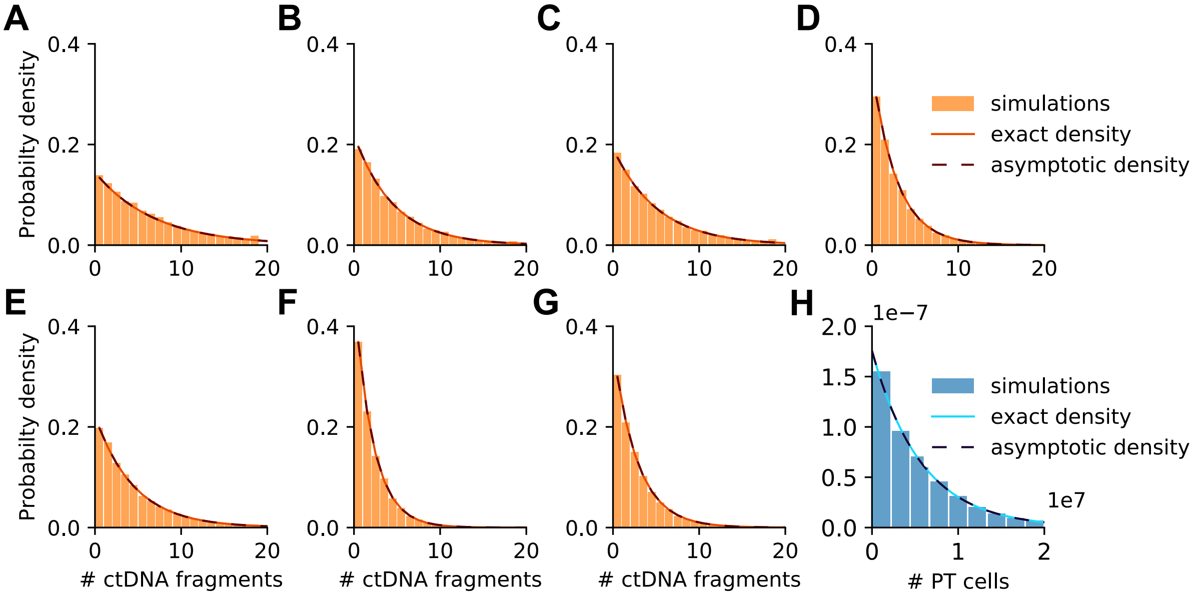


**Fig. S3: Comparison between computer simulations, exact and asymptotic theoretical distributions conditioned on tumor survival.** Panels (**A**–**G**) show the level of biomarker shed by cancer cells and circulating in the bloodstream at time $t$. Bars illustrate the distribution of the number of biomarker molecules based on $10^4$ simulation realizations. The probability distributions illustrated are all conditional on primary tumor survival up to time $t$. Full lines illustrate the exact probability distribution at time $t$, while dashed lines represent the asymptotic results for large times and small shedding rate. Both agree with simulation results (bars). Each panel shows the following combinations of shedding rates: **A** | $q_b > 0, \lambda_1 > 0, q_d > 0$. **B** | $q_b > 0, \lambda_1 > 0, q_d = 0$. **C** | $q_b > 0, \lambda_1 = 0, q_d > 0$. **D** | $q_b > 0, \lambda_1 = 0, q_d = 0$. **E** | $q_b = 0, \lambda_1 > 0, q_d > 0$. **F** | $q_b = 0, \lambda_1 > 0, q_d = 0$. **G** | $q_b = 0, \lambda_1 = 0, q_d > 0$. **H** | Probability distribution of the primary tumor size at a given time $t$. Parameter values: birth rate $b = 0.14$ per cell per day and death rate $d = 0.136$ per cell per day; ctDNA half-life time $t_{1/2} = 30$ minutes; ctDNA shedding probability per cell death $q_d = 10^{-4}$, per cell division $q_b = 10^{-4}$, and shedding rate per day (at cell necrosis) $\lambda_1 = 10^{-5}$ per cell per day. All results are computed at time $t = 3000$ days from the primary tumor onset.

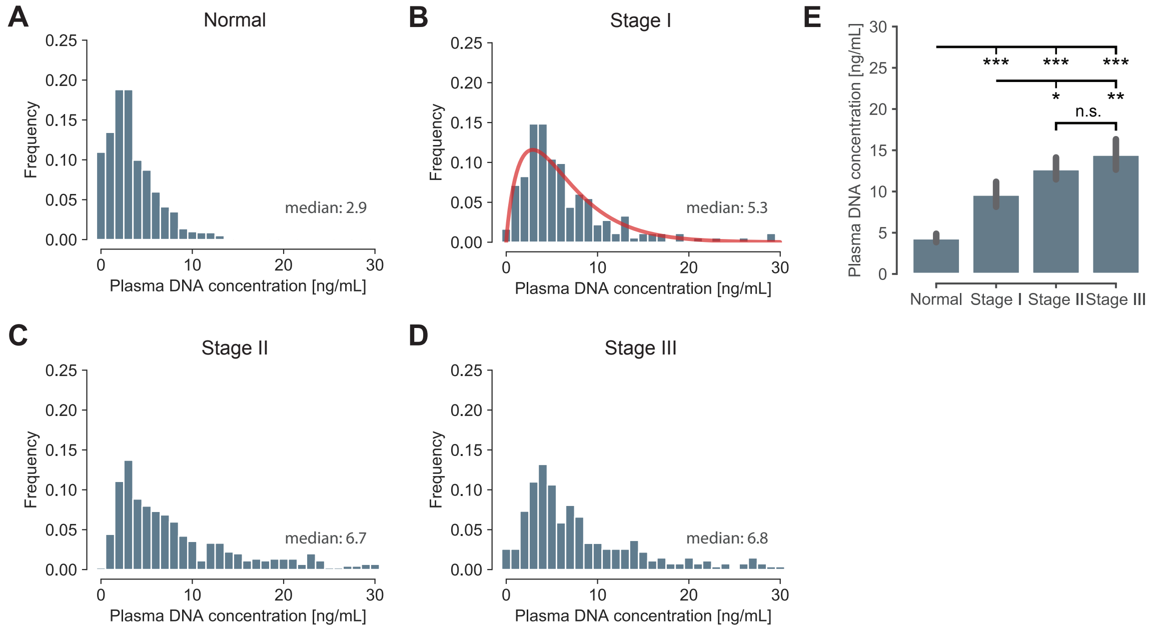


**Fig. S4: Plasma DNA concentration increases with disease progression.** Analyzed data from 812 normal subjects, 199 stage I patients, 497 stage II patients, and 307 stage III patients of Cohen et al (*9*). **A**−**D** | Plasma DNA concentrations (ng per plasma mL) in healthy and cancer patients. **b** | Red line illustrates fitted Gamma distribution with mean of 6.3 and median of 5.2 ng per plasma mL. Depicted distribution is used to model the varying plasma DNA concentration. **E** | Comparing plasma DNA concentrations. Two-sided Wilcoxon rank-sum tests were used. Thick black bars denote 90% confidence interval. * $P < 0.05$; ** $P < 0.01$; *** $P < 0.001$; n.s. denotes not significant.

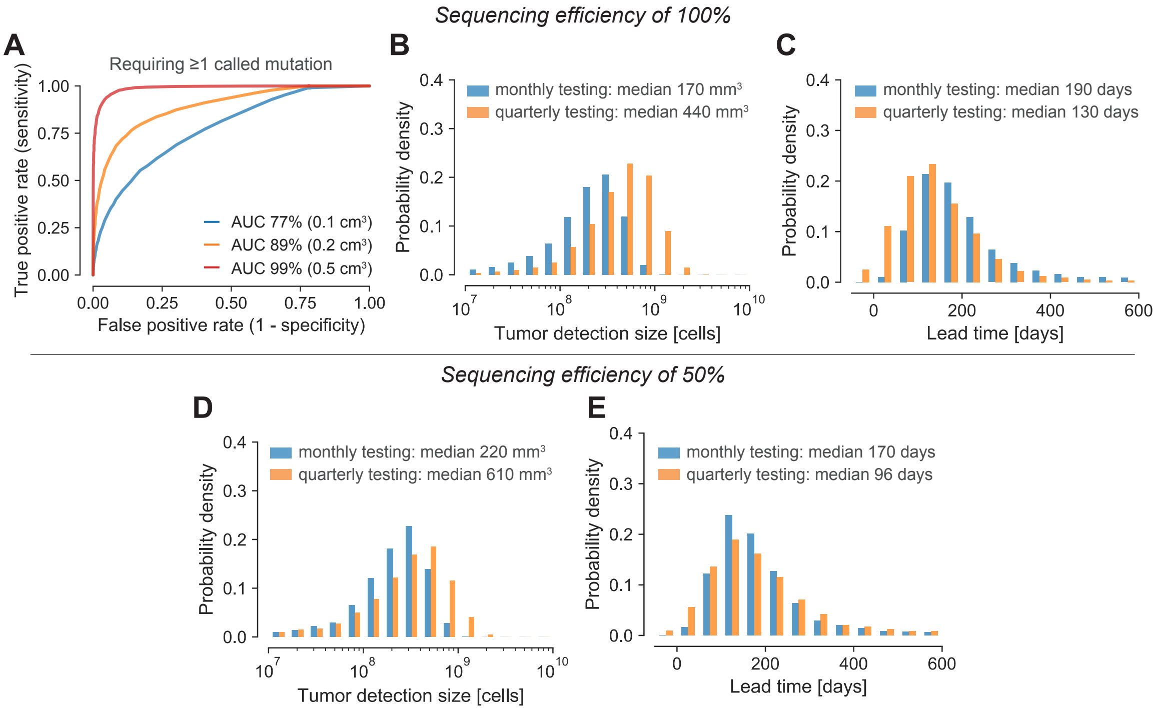


**Fig. S5: Expected tumor detection size and lead time compared to current clinical relapse detection. A** | ROC (Receiver Operating Characteristic) curves for tumors with 100 million cells ($\approx 0.1$ cm$^3$, blue line), 200 million cells ($\approx 0.2$ cm$^3$, orange line), or 500 million cells ($\approx 0.5$ cm$^3$, red line) when 20 clonal tumor specific mutations are tracked for relapse detection and one of these 20 mutations needs to be called for a positive test. AUC, area under the curve. **B–E** | Since the same relapse detection test with a specificity of 99.5% for monthly and quarterly testing was used, the monthly sampling produces 0.06 false-positives over 12 months of relapse testing while the quarterly sampling only produces 0.02 false-positives over 12 months. **B** | Expected tumor detection size distributions for monthly and quarterly repeated relapse detection with a fixed specificity of 99.5% per test and a sequencing efficiency of 100%. **C** | Expected lead time distributions compared to imaging-based approaches applied at the same frequency with a detection limit of 1 cm$^3$ for monthly and quarterly repeated relapse detection with a fixed specificity of 99.5% per test and a sequencing efficiency of 100%. **D** | Expected tumor detection size distributions for monthly and quarterly repeated relapse detection with a fixed specificity of 99.5% per test and a sequencing efficiency of 50%. **E** | Expected lead time distributions compared to imaging-based approaches applied at the same frequency with a detection limit of 1 cm$^3$ for monthly and quarterly repeated relapse detection with a fixed specificity of 99.5% per test and a sequencing efficiency of 50%. Parameter values typical for lung cancer: birth rate $b = 0.14$ per cell per day; death rate $d = 0.13$ per cell per day; ctDNA half-life time $t_{1/2} = 30$ minutes; ctDNA shedding probability per cell death $q_d = 1.4 \cdot 10^{-4}$; sequencing panel covers 20 clonal tumor-specific mutations; sequencing error rate per base-pair $1.5 \cdot 10^{-5}$; 15 mL blood sampled per test; plasma DNA median concentration 5.2 ng per plasma mL.

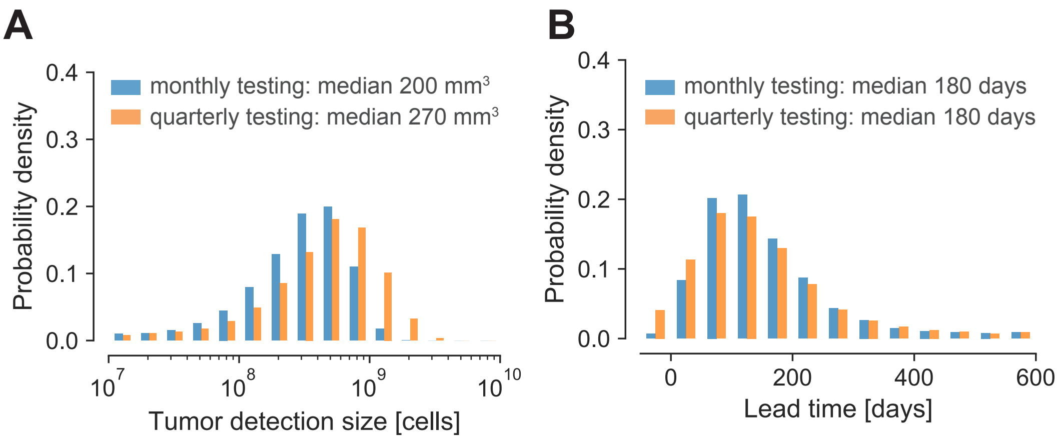


**Fig. S6: Expected tumor detection size and lead time compared to current clinical relapse detection for a sequencing efficiency of 100%.** For better comparability, positive detection test thresholds were set such that if the test is repeated multiple times, a combined false positive rate of 5% is obtained over all tests per year. Note the decreased expected detection sizes and increased lead times compared to Fig. 3 where a sequencing efficiency of 50% was assumed. **A** | Expected tumor detection size distributions for monthly and quarterly repeated relapse detection tests. **B** | Expected lead time distributions compared to imaging-based approaches applied at the same frequency with a detection limit of 1 $cm^3$ for monthly and quarterly repeated relapse detection tests. Parameter values typical for lung cancer: birth rate $b = 0.14$ per cell per day; death rate $d = 0.13$ per cell per day; ctDNA half-life time $t_{1/2} = 30$ minutes; ctDNA shedding probability per cell death $q_d = 1.4 \cdot 10^{-4}$; sequencing panel covers 20 clonal tumor-specific mutations; sequencing error rate per base-pair $1.5 \cdot 10^{-5}$; sequencing efficiency 100%; 15 mL blood sampled per test; plasma DNA median concentration 5.2 ng per plasma mL.

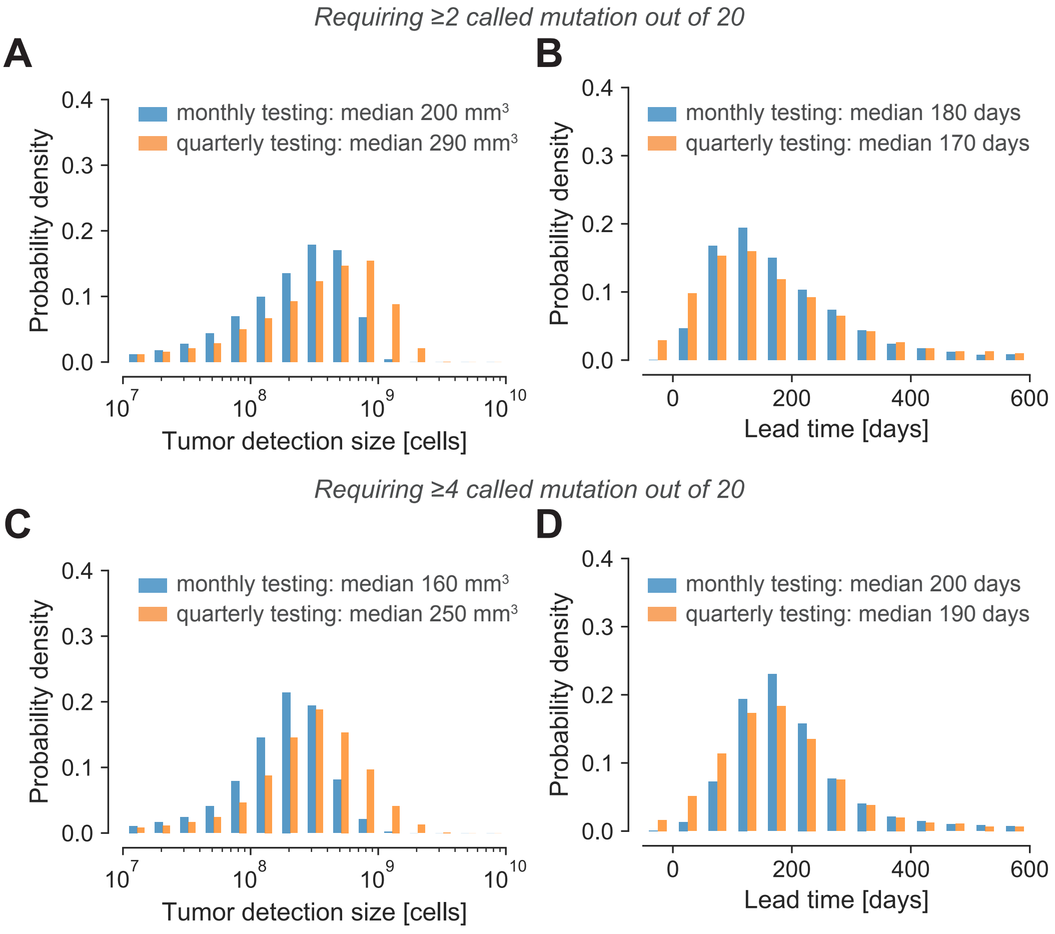


**Fig. S7: Expected tumor relapse detection size and lead time compared to current clinical relapse detection when multiple called mutations are required for a positive test.** **A** | Expected tumor detection size distributions for monthly and quarterly repeated relapse detection tests when 20 a priori known mutations are covered by the sequencing panel and two of these 20 mutations need to be called for a positive relapse detection. **B** | Expected lead time distributions compared to imaging-based approaches applied at the same frequency with a detection limit of 1 $cm^3$ for monthly and quarterly repeated relapse detection tests. **C** | Expected tumor detection size distributions for monthly and quarterly repeated relapse detection tests when 20 a priori known mutations are covered by the sequencing panel and four of these 20 mutations need to be called for a positive relapse detection. **D** | Expected lead time distributions compared to imaging-based approaches applied at the same frequency with a detection limit of 1 $cm^3$ for monthly and quarterly repeated relapse detection tests. Parameter values typical for lung cancer: birth rate $b = 0.14$ per cell per day; death rate $d = 0.13$ per cell per day; ctDNA half-life time $t_{1/2} = 30$ minutes; ctDNA shedding probability per cell death $q_d = 1.4 \cdot 10^{-4}$; sequencing panel covers 20 clonal tumor-specific mutations; sequencing error rate per base-pair $1.5 \cdot 10^{-5}$; sequencing efficiency 50%; 15 mL blood sampled per test; plasma DNA median concentration 5.2 ng per plasma mL. For better comparability, positive detection test thresholds were set such that if the test is repeated multiple times, a combined false positive rate of 5% is obtained over all tests per year.

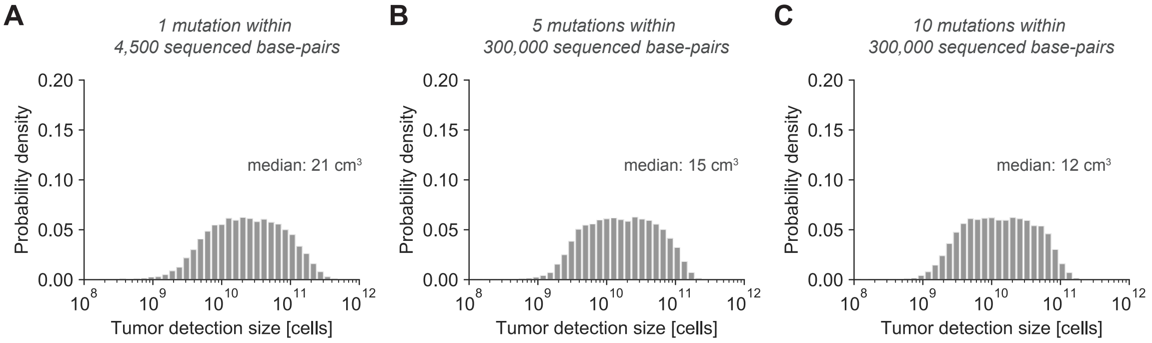


**Fig. S8: Expected tumor detection size distributions for fast growing lung cancers.** Tumors grow with a rate of $r = 1.0\%$ per day which corresponds to a tumor volume doubling time of 70 days. Positive screening test thresholds were set such that 0.01 false-positives are expected per year (i.e., specificity of 99% for an annually repeated test). **A** | Repeated virtual screening tests with one expected mutation within a sequencing panel covering 4,500 base-pairs. **B** | Repeated virtual screening tests with five expected mutation within a sequencing panel covering 300,000 base-pairs. **C** | Repeated virtual screening tests with ten expected mutation within a sequencing panel covering 300,000 base-pairs. Parameter values: birth rate $b = 0.14$ per cell per day; death rate $d = 0.13$ per cell per day; ctDNA half-life time $t_{1/2} = 30$ minutes; ctDNA shedding probability per cell death $q_d = 1.4 \cdot 10^{-4}$; sequencing error rate per base-pair $1.5 \cdot 10^{-5}$; sequencing efficiency 50%; 15 mL blood sampled per test; plasma DNA median concentration 5.2 ng per plasma mL.

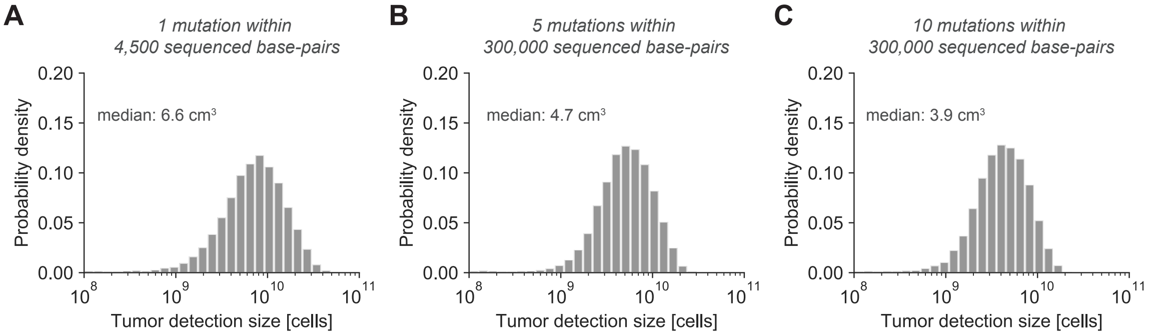


**Fig. S9: Expected tumor detection size distributions in the presense of eight lung nodules.** Positive screening test thresholds were set such that 0.01 false-positives are expected per year of screening (i.e., specificity of 99% for an annually repeated test). Assuming eight benign lung nodules with each 34 million cells (diameter of $\approx$ 4 mm) replicating at a constant size with a cell birth/death rate of $b_{\text{bn}} = d_{\text{bn}} = 0.07$ per day and shedding ctDNA with the same probability $q_d$ (*26, 46*). Sequencing panels were conservatively assumed to cover the same number of somatic mutations in benign nodules and malignant tumors. Repeated virtual screening tests with one mutation per tumor covered by a sequencing panel of 4,500 base-pairs (panel **A**), with five mutations per tumor (panel **B**) and with ten mutations per tumor covered by a sequencing panel of 300,000 base-pairs (panel **C**). Parameter values: birth rate $b = 0.14$ per cell per day; death rate $d = 0.136$ per cell per day; ctDNA half-life time $t_{1/2} = 30$ minutes; ctDNA shedding probability per cell death $q_d = 1.4 \cdot 10^{-4}$; sequencing error rate per base-pair $1.5 \cdot 10^{-5}$; sequencing efficiency 50%; 15 mL blood sampled per test; plasma DNA median concentration 5.2 ng per plasma mL.

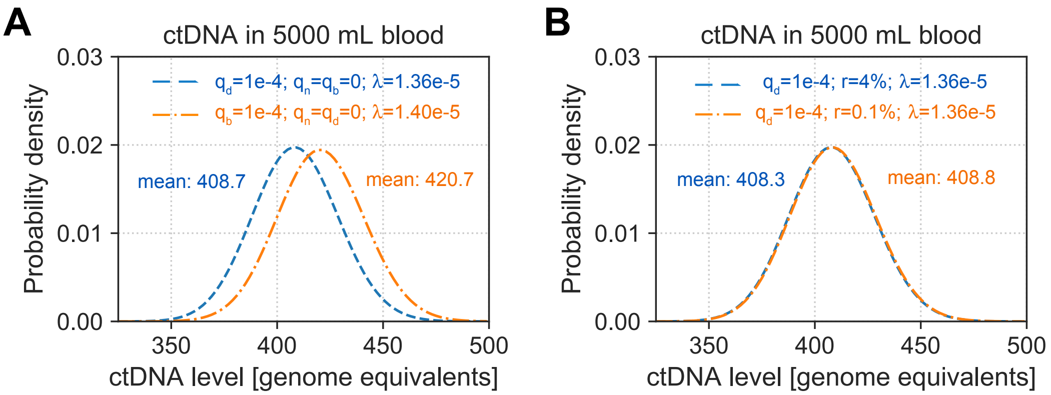


**Fig. S10: Operating ctDNA shedding processes determine the effective shedding rate.** Panels depict diploid genome equivalents of ctDNA present in 5 liters of blood (2.75 liters of plasma) when a lung tumor reaches a a size of 1 billion cells. **A** | Blue line illustrates a scenario where ctDNA is exclusively shed during apoptosis with a ctDNA shedding probability per cell death $q_d = 10^{-4}$. Orange line illustrates a scenario where ctDNA is exclusively shed during proliferation with a ctDNA shedding probability per cell division $q_b = 10^{-4}$. We obtain a shedding rate of $\lambda = 1.36 \cdot 10^{-5}$ per day per cell for the first case and a shedding rate of $\lambda = 1.40 \cdot 10^{-5}$ for the second case because cells proliferate and shed ctDNA slightly more frequently than they die and shed ctDNA ($b = 0.14 > d = 0.136$). **B** | In both scenarios cells shed ctDNA exclusively during apoptosis ($q_d = 10^{-4}$, $q_n = q_b = 0$). Cells die with the same death rate $d = 0.136$ per cell per day and hence, we obtain identical shedding rates of $\lambda = 1.36 \cdot 10^{-5}$ per cell per day. In the blue scenario, cells divide with a rate of $b = 0.137$ per day while cells divide with a rate of $b = 0.176$ per day in the orange scenario. Although the shedding rates are identical, the slow growth rate of $r = 0.1\%$ leads to a slightly higher ctDNA amount at the same size of a tumor. Parameter values: ctDNA half-life time $t_{1/2} = 30$ minutes.

#### Supplementary Tables

**Table S1: Mean and variance for the distribution of the circulating biomarker $C_t^A$ at time $t$ shed by a tumor conditioned on survival until time $t$.** Simulated results are averages over $10^4$ realizations and are compared with the theoretical expression derived from the marginal probability generating function $\mathscr{C}_t^A(y \mid \Omega_t)$. The first column shows the seven combinations of shedding rates we considered. Parameter values: birth rate $b = 0.14$ per cell per day, death rate $d = 0.136$ per cell per day, ctDNA half-life time $t_{1/2} = 30$ minutes, tumor age $t = 3500$ days corresponding to an expected size of $4.209 \times 10^7$ cells.

| **Shedding dynamics** | | | **Mean** | | | **Variance** | | |
|---|---|---|---|---|---|---|---|---|
| $\lambda_0$ | $\lambda_1$ | $\lambda_2$ | Simulations | Exact | Asymptotic | Simulations | Exact | Asymptotic |
| 0 | $10^{-5}$ | 0 | 12.5298 | 12.6494 | 12.6494 | 169.1204 | 172.6576 | 172.6579 |
| $1.36 \times 10^{-5}$ | 0 | 0 | 17.1683 | 17.2032 | 17.2032 | 305.7878 | 313.1543 | 313.1548 |
| 0 | 0 | $1.4 \times 10^{-5}$ | 17.5059 | 17.7092 | 17.7092 | 326.4480 | 331.3252 | 331.3257 |
| $1.36 \times 10^{-5}$ | $10^{-5}$ | 0 | 30.1496 | 29.8527 | 29.8527 | 940.0496 | 921.0341 | 921.0356 |
| 0 | $10^{-5}$ | $1.4 \times 10^{-5}$ | 30.4422 | 30.3586 | 30.3587 | 956.3959 | 952.0057 | 952.0072 |
| $1.36 \times 10^{-5}$ | 0 | $1.4 \times 10^{-5}$ | 34.4846 | 34.9124 | 34.9125 | 1235.4580 | 1253.7906 | 1254.7926 |
| $1.36 \times 10^{-5}$ | $10^{-5}$ | $1.4 \times 10^{-5}$ | 47.3219 | 47.5619 | 47.5619 | 2284.9660 | 2309.6932 | 2309.6970 |

**Table S2: Mean and variance for the distribution of the circulating biomarker $C_t^A$ at time $t$ shed by a tumor conditioned on survival until time $t$ across a wide range of shedding rates.** Simulated results are averages over $10^5$ realizations and are compared with the theoretical expression derived from the marginal probability generating function $\mathscr{C}_t^A(y \mid \Omega_t)$. We explore a wide range of shedding rates by varying $\lambda_0$ and setting $\lambda_1 = \lambda_2 = 0$. Parameter values: birth rate $b = 0.14$ per cell per day, death rate $d = 0.136$ per cell per day, ctDNA half-life time $t_{1/2} = 30$ minutes, tumor age $t = 3000$ days corresponding to an expected size of $5.696 \times 10^6$ cells.

| **Shedding dynamics** | **Mean** | | | **Variance** | | |
|---|---|---|---|---|---|---|
| $\lambda = \lambda_0$ | Simulations | Exact | Asymptotic | Simulations | Exact | Asymptotic |
| $10^{-3}$ | 171.6 | 171.04203 | 171.04305 | 29201 | 29426.407 | 29426.768 |
| $10^{-4}$ | 17.13 | 17.104203 | 17.104305 | 306.8 | 309.65785 | 309.66155 |
| $10^{-5}$ | 1.713 | 1.7104203 | 1.7104305 | 4.614 | 4.6359567 | 4.6360029 |
| $10^{-6}$ | 0.1709 | 0.1710420 | 0.1710430 | 0.2005 | 0.2002974 | 0.2002988 |
| $10^{-7}$ | 0.0172 | 0.0171042 | 0.0171043 | 0.01751 | 0.0173968 | 0.0173969 |
| $10^{-8}$ | 0.001755 | 0.0017104 | 0.0017104 | 0.001756 | 0.0017133 | 0.0017134 |

**Table S3: Mean number of mutations of lung cancers covered by CancerSEEK and CAPP-Seq.** Mean number of clonal mutations of early stage lung cancers in the TRACERx study (*27*) that are covered by the CAPP-Seq panel and the CancerSEEK panel (considing all amplicons entirely).

| Cohort | Filter | CAPP-Seq | CancerSEEK |
|---|---|---|---|
| Whole | Overall | 9.14 | 1.12 |
| | Never Smoker | 3.83 | 1.00 |
| | Ex Smoker | 8.25 | 1.02 |
| | Recent Ex Smoker | 12.12 | 1.33 |
| | Current Smoker | 10.29 | 1.00 |
| Stage I Only | Overall | 9.71 | 1.03 |
| | Never Smoker | 4.67 | 1.22 |
| | Ex Smoker | 8.76 | 0.83 |
| | Recent Ex Smoker | 12.80 | 1.30 |
| | Current Smoker | 12.50 | 0.75 |

#### Supplementary Note S1: Mathematical Modeling

We developed a stochastic model of cancer evolution and biomarker shedding to study the potential of cancer detection based on the biomarker level in liquid biopsies. Here we focused on somatic point mutations present in circulating tumor DNA (ctDNA). Nonetheless, our framework applies to any biomarker that is shed by cancer cells at some rate. These biomarkers can be released at different stages of a cell cycle, for example during proliferation, apoptosis, or per unit of time.

We model the dynamics of tumor growth and biomarker shedding through a continuous-time multi-type branching process (*31–33, 59, 60*). The primary tumor grows stochastically from a single malignant cell at time $t = 0$. The tumor size at time $t$, denoted $A_t$ and expressed in number of cells, is modeled by a supercritical branching birth death process with net growth rate $r = b - d > 0$, where $b$ and $d$ denote the birth and death rate per day, respectively (Fig. 1B). This process has a probability of $\delta = d/b$ to go extinct (*59*). Since we are only interested in primary tumors that do not go extinct, we condition $A_t$ on eventual survival, i.e. on the event $\Omega := \{A_t > 0 \text{ for all } t\}$. Cancer cells shed a biomarker into the bloodstream at a rate $\lambda$ per day. In our case, the biomarker is ctDNA measured in diploid genome equivalents (GE). For now, we assume that the biomarker is exclusively shed by cells undergoing apoptosis and hence the shedding rate is given by $\lambda = d \cdot q_d$, where $q_d$ denotes the shedding probability per cell death (Fig. 1B).

Normal or benign tumor cells also shed cell-free DNA (cfDNA) into the bloodstream. Cells in benign tumors and expanded subclones often harbor the same cancer-associated mutations as cancer cells (*42–44, 61, 62*). Hence, ctDNA shed from benign tumors can be difficult to distinguish from ctDNA shed from malignant tumors. We define $B_t$ as the size of the benign population of cells at time $t$ that shed the biomarker in the bloodstream, and denote their shedding rate as $\lambda_{\text{bn}}$ . We assume that benign lesions roughly replicate at a constant size. Hence, benign cells divide and die at the same rate $b_{\text{bn}} = d_{\text{bn}}$, and their population size $B_t$ remains constant over time ($B_t = B_0$ for all $t$). Considering again that the biomarker is exclusively shed by cells undergoing apoptosis, the shedding rate of benign cells is $\lambda_{\text{bn}} = d_{\text{bn}} \cdot q_{d,\text{bn}}$ per day. Although we conservatively assume that apoptotic benign cells shed ctDNA with the same probability as malignant cells ($q_d = q_{d,\text{bn}}$), the shedding rate $\lambda_{\text{bn}}$ of benign cells is lower than the shedding rate $\lambda$ of malignant cells because benign cells typically replicate at a lower rate than cancer cells (*26*).

The circulating biomarker is eliminated from the bloodstream at an elimination rate $\varepsilon$ which can be calculated from the biomarker half-life time $t_{1/2}$ as $\varepsilon = \log(2)/t_{1/2}$. We denote as $C_t^A$ and $C_t^B$ the amount of biomarker circulating in the bloodstream at time $t$ shed by malignant and benign cells, respectively. The total amount of the biomarker circulating at time $t$ is thus $C_t = C_t^A + C_t^B$.

Because malignant and benign cells shed the biomarker independently from each other, the processes $(A_t, C_t^A)$ and $(B_t, C_t^B)$ can be studied separately. The stochastic process $(A_t, C_t^A)$ is a two-type branching process governed by the following transitions

$$\begin{aligned} A &\longrightarrow AA && \text{rate } b \\ A &\longrightarrow C && \text{rate } d \cdot q_d \\ A &\longrightarrow \emptyset && \text{rate } d\,(1-q_d) \\ C &\longrightarrow \emptyset && \text{rate } \varepsilon\ . \end{aligned}$$

The process is initialized at time $t = 0$ with a single cancer cell and no circulating biomarkers, that is $\left(A_0, C_0^A\right) = (1, 0)$.

Because we assume that the benign cell population $B_t$ remains constant over time and biomarker units are eliminated from the bloodstream independently at the same rate, the process $C_t^B$ is a branching pure-death process with constant immigration (*63*) (or equivalently a $M/M/\infty$ queue system (*64, 65*)). We assume that $C_t^B$ is at equilibrium at time $t = 0$. We present additional details about this process, including the exact form of $C_0^B$, in the next section.

**Biomarker Size Distributions**

The processes $A_t$, $B_t$ and $C_t$ are indexed by the time parameter $t$, counted from the primary tumor onset. Clinically, this time is impossible to measure, and it is thus convenient to express the previous processes in terms of other indexing parameters. In our case, a suitable choice is to use the first passage time when the tumor reaches a size of $m$ cells

$$\tau_m := \inf\{t > 0 : A_t = m\}\ .$$

By employing this random variable, the processes $C_{\tau_m}^A$ and $C_{\tau_m}^B$ express the biomarker levels in terms of an observable quantity, i.e. the primary tumor size. We are especially interested in our model predictions for relatively large primary tumor sizes. When $m$ is large and the shedding rate $\lambda$ is small, we are able to apply asymptotic results for the distributions of the processes $C_{\tau_m}^A$ and $C_{\tau_m}^B$ that greatly simplify computations and help getting insights into the main evolutionary features of the model. Hence, we first focus on these asymptotic results, while later we will show how to generalize our framework to include multiple biomarker shedding dynamics and to derive the probability distribution of the processes $A_t$, $B_t$ and $C_t$ indexed by time. Unless otherwise specified, the presented results are conditioned on eventual survival of $A_t$.

Note that we will employ the same sign '$\sim$' to indicate "distributed as" and "asymptotically converges to". Since only the former usage applies to random variables, this does not cause any notational ambiguity. Finally, whenever asymptotic probability distributions are derived, formulas including the '$\sim$' sign will be accompanied by text highlighting the relevant asymptotic limit.

##### *Process* $C_t^A$

Here we provide a sketch derivation for the asymptotic distribution of $C_{\tau_m}^A$ in the small shedding rate - large primary tumor size limit ($\lambda \ll 1$, $m \gg 1$). The derivation follows from the results presented in ref. (*66*).

The biomarker is shed by cancer cells as a Cox process (or doubly stochastic Poisson process (*67*)) with intensity $(\lambda \cdot A_t)_{t \geq 0}$. Furthermore, for large times the branching birth death process $A_t$ satisfies the classic result (*59*)

$$\lim_{t \to \infty} A_t = W e^{rt} \tag{1}$$

where $W$ is a non-negative random variable such that $W = 0$ if and only if the process $A_t$ goes extinct. Let us assume $m \gg 1$ and consider the process $A_t$ at time $\tau_m$, by definition, that is when the primary tumor is made of $m$ cells. Conditioning on $\Omega \equiv \{W > 0\}$ and looking backwards in time, we see that the size of the primary tumor at $s$ days before $\tau_m$ is given by $m \cdot e^{-rs}$. Hence, in the small $\lambda$ - large $m$ limit, the biomarker is shed as a Poisson process with mean $\lambda \int_0^\infty m e^{-rs} ds = \lambda \cdot m/r$. Moreover, by the same argument we see that the unordered times of biomarker shedding are asymptotically distributed as i.i.d. Exponential($r$) random variables. Given that the elimination times of shed biomarkers are also i.i.d. Exponential($\varepsilon$), the probability that a randomly selected biomarker fragment is still circulating in the bloodstream at time $\tau_m$ is $r/(\varepsilon + r)$. Due to the thinning property of Poisson processes, we find that for large primary tumor sizes $m$ and small shedding rates $\lambda$

$$C_{\tau_m}^{A} \sim \text{Poisson}\left(\frac{m \cdot \lambda}{\varepsilon + r}\right) \ . \tag{2}$$

##### ***Process*** $C_t^B$

The process $C_t^B$ is a branching pure-death process with immigration, with death rate $\varepsilon$ and immigration rate $B_0 \cdot \lambda_{\text{bn}} = B_0 \cdot d_{\text{bn}} \cdot q_d$, because the population size $B_t$ is constant over time. The probability generating function for such a process follows from ref. (*63*)

$$\mathscr{C}^B(y,t) = \left(1 + (y-1)e^{-\varepsilon t}\right)^{C_0^B} e^{\frac{\lambda_{\text{bn}}}{\varepsilon} B_0 (y-1)(1-e^{-\varepsilon t})} \ . \tag{3}$$

Conditioning on $A_t$ survival, we have that $\lim_{m\to\infty} \tau_m = \infty$ almost surely. Therefore, since the process $(B_t, C_t^B)$ is independent of $A_t$, the large $m$ limit of $C_{\tau_m}^B$ coincides with the large time limit of $C_t^B$. When $t$ is large, equation (3) converges to

$$\lim_{t\to\infty} \mathscr{C}^B(y,t) = e^{\frac{\lambda_{\text{bn}}}{\varepsilon} B_0 (y-1)} \ . \tag{4}$$

independently of the initial biomarker level. The right hand side of equation (4) is the probability generating function of a Poisson random variable with mean $B_0 \cdot \lambda_{\text{bn}}/\varepsilon$, and so for large $m$ we have

$$C_{\tau_m}^{B} \sim \text{Poisson}\left(\frac{B_0 \cdot \lambda_{\text{bn}}}{\varepsilon}\right) \ . \tag{5}$$

Since we assume that $C_t^B$ is in equilibrium initially, we set $C_0^B = B_0 \cdot d_{\text{bn}} \cdot q_d/\varepsilon$.

##### ***Process*** $C_t$ ***and sampling***

By combining the previous results, we find the asymptotic limit for the distribution of the total biomarker amount present in the bloodstream when the primary tumor is made of a large number of cells. Conditioned on $A_t$ survival, this limit is given by

$$C_{\tau_m} = C_{\tau_m}^{A} + C_{\tau_m}^{B} \xrightarrow{d} \text{Poisson}\left(\frac{m\, d\, q_d}{\varepsilon + r} + \frac{B_0\, d_{\text{bn}}\, q_d}{\varepsilon}\right) \tag{6}$$

Suppose that a sample of blood with volume $V_s$ is drawn from a patient with a total blood volume equal to $V_{tot}$. Assuming that the circulating biomarker units are well mixed in the bloodstream, we sample the biomarker units from the bloodstream with probability $p = V_s/V_{tot}$. We define $X_t$ as the total biomarker amount (i.e., number of diploid genome equivalents) present in a sample drawn at time $t$. Then, by applying again the thinning property of Poisson processes, we find that for large $m$ the biomarker amount in the sample when the primary tumor is made of $m$ cells is asymptotically distributed as

$$X_{\tau_m} \sim \text{Poisson}\left(p\left[\frac{m\,d\,q_d}{\varepsilon + r} + \frac{B_0\,d_{\text{bn}}\,q_d}{\varepsilon}\right]\right)\ . \tag{7}$$

**General Framework**

For the stochastic analysis of ctDNA shedding by malignant and benign cells we employed the asymptotic results derived above. We further extended these results in several ways. First, so far we focused on the case where biomarker shedding occurs exclusively at cell apoptosis. We generalize our derivations to additionally include shedding at cell necrosis (per unit of time) and proliferation. Second, we presented size distributions in the asymptotic limit of large primary tumor sizes - small shedding rates. For other shedding processes, We obtain similar distributions in the asymptotic limit for large times - small shedding rates. Finally, we compute exact probability distributions for the processes $A_t$, $B_t$ and $C_t$ at any given time $t$ for all three shedding processes and their mixtures. Below we present these three extensions in greater detail.

In order to include the possibility of biomarker shedding by cancerous cells also at necrosis and proliferations in our model, we consider the following set of transitions for the process $(A_t, C_t)$

$$\begin{array}{lll}
A \longrightarrow AAC & \text{rate } b\,q_b & \text{rate } \lambda_2 \\
A \longrightarrow AA & \text{rate } b\,(1-q_b) & \text{rate } b-\lambda_2 \\
A \longrightarrow AC & \text{rate } q_n & \text{rate } \lambda_1 \\
A \longrightarrow C & \text{rate } d\,q_d & \text{rate } \lambda_0 \\
A \longrightarrow \emptyset & \text{rate } d\,(1-q_d) & \text{rate } d-\lambda_0 \\
C \longrightarrow \emptyset & \text{rate } \varepsilon & \text{rate } \varepsilon\ .
\end{array} \tag{8}$$

Here, $q_d$ and $q_b$ denote the shedding probabilities at apoptosis and proliferation, respectively. Similarly, $\lambda_0 = d\,q_d$, $\lambda_1 = q_n$ and $\lambda_2 = b\,q_b$ denote the shedding rates of malignant cells at apoptosis, necrosis, and proliferation, respectively. The total shedding rate is then defined by $\lambda = \lambda_0 + \lambda_1 + \lambda_2$.

***Asymptotic results for multiple shedding processes***

With the generalized definition of the total shedding rate $\lambda$, we find that the biomarker is shed by cancer cells as a Cox process with intensity $(\lambda A_t)_{t\geq 0}$. Hence, the derivation of $C^A_{\tau_m}$ distribution is identical to the one above. In particular, we show again that for a large primary tumor size $m$ and a small total shedding rate $\lambda$, the biomarker amount present in the bloodstream when $A_t = m$ asymptotically follows a Poisson distribution with mean $\frac{\lambda \cdot m}{\varepsilon + r}$.

Asymptotic results for large times

A similar derivation provides the size distribution for $C^A_t$ in the asymptotic limit for large time and small total shedding rate. Under this limit and conditioning on $A_t$ survival, the total biomarker amount shed up to time $t$ converges in finite dimensional distributions to a Poisson random variable with mean $\frac{W\lambda e^{rt}}{r}$, where $W$ is the same as in equation (1) (for details see ref. (*66*)). We recall that $W > 0$ if and only if $A_t$ eventually survives, and conditioned on this event, $W$ follows an Exponential $\left(\frac{r}{b}\right)$ distribution. The biomarker amount still present in the bloodstream at time $t$ is thus a compound Poisson process with probability generating function

$$\mathrm{E}\left[z^{C^A_t}\right] = \mathrm{E}\left[\exp\left(\frac{W\lambda e^{rt}}{r}\left(\mu(z) - 1\right)\right)\right] .$$

Here $\mu(z)$ denotes the probability generating function of the process initiated by a randomly selected biomarker unit, but because the biomarker cannot reproduce, this process is a branching pure-death process whose size can only be 1 or 0. The probability generating function of such a process is thus $\mu(z) = \frac{\varepsilon + rz}{\varepsilon + r}$. Using this expression and the results for $W$, the probability generating function for $C^A_t$ becomes

$$\mathrm{E}\left[z^{C^A_t}\right] = \left(1 - \frac{b\lambda e^{rt}(z-1)}{r(\varepsilon + r)}\right)^{-1} . \tag{9}$$

Notice that the same steps can be repeated to formally derive the asymptotic results for large primary tumor sizes. The last expression represents the probability generating function of a geometric random variable, and so in the asymptotic limit for large times and small total shedding rates, conditioned on $A_t$ survival, we find that

$$C_t^A \sim \text{Geometric}\left(\frac{r(\varepsilon+r)}{b\lambda e^{rt}+r(\varepsilon+r)}\right) \ . \tag{10}$$

*First moments*

In this section, we summarize the expected values and variances of the processes $A_t$ and $C_t$ in the considered asymptotic limits. These moments follow from the probability distributions we previously derived. The process $A_t$ is a supercritical branching birth death process with net growth rate $r$ and extinction probability $\delta = d/b$. In particular, recalling that $\Omega$ denotes the event of $A_t$ eventual survival, we have $\mathrm{P}(\Omega) = 1-\delta$. If $A_0 = 1$, we have

$$\mathrm{E}[A_t] = e^{rt}\,, \quad \mathrm{Var}(A_t) \sim \frac{1+\delta}{1-\delta}\, e^{2rt}$$

and

$$\mathrm{E}[A_t \mid \Omega\,] \sim \frac{e^{rt}}{1-\delta}\,, \quad \mathrm{Var}(A_t \mid \Omega\,) \sim \frac{1+2\delta}{(1-\delta)^2}\, e^{2rt} \ .$$

For the process $C_t^A$, given that $(A_0, C_0^A) = (1,0)$, equations (2) and (10) yield

$$\mathrm{E}\left[C_{\tau_m}^A \mid \Omega\,\right] \sim \frac{m\ \lambda}{\varepsilon+r}, \quad \mathrm{Var}\left(C_{\tau_m}^A \mid \Omega\,\right) \sim \frac{m\ \lambda}{\varepsilon+r} \tag{11}$$

and

$$\mathrm{E}\left[C_t^A \mid \Omega\,\right] \sim \frac{b\lambda}{r(\varepsilon+r)}\, e^{rt}\,, \quad \mathrm{Var}\left(C_t^A \mid \Omega\,\right) \sim \frac{b\lambda}{r(\varepsilon+r)}\, e^{rt}\left(\frac{b\lambda}{r(\varepsilon+r)}\, e^{rt}+1\right) \tag{12}$$

asymptotically for large primary tumor sizes - small shedding rates and large times - small shedding rates, respectively.

In the same two limits, the process $C_t^B$ exhibits the same asypmtotic behaviour. Conditional on $C_0^B = B_0 \cdot \lambda_{\mathrm{bn}}/\varepsilon$ we find

$$\lim_{m\to\infty} \mathrm{E}\left[C_{\tau_m}^B\right] = \lim_{t\to\infty} \mathrm{E}[C_t^B] = \frac{\lambda_{\mathrm{bn}}}{\varepsilon} B_0\,, \quad \lim_{m\to\infty} \mathrm{Var}\left(C_{\tau_m}^B\right) = \lim_{t\to\infty} \mathrm{Var}\left(C_t^B\right) = \frac{\lambda_{\mathrm{bn}}}{\varepsilon} B_0 \ .$$

Combining these results and applying the same initial conditions, we then get

$$\mathrm{E}\left[C_{\tau_m} \mid \Omega\,\right] \sim \frac{m\ \lambda}{\varepsilon+r} + \frac{B_0\ \lambda_{\mathrm{bn}}}{\varepsilon}, \quad \mathrm{Var}\left(C_{\tau_m} \mid \Omega\,\right) \sim \frac{m\ \lambda}{\varepsilon+r} + \frac{B_0\ \lambda_{\mathrm{bn}}}{\varepsilon} \tag{13}$$

for large $M$ - small $\lambda$ and

$$\begin{aligned} \mathrm{E}\left[C_t \mid \Omega\right] &\sim \frac{b\ \lambda}{r(\varepsilon+r)}\, e^{rt} + \frac{B_0\ \lambda_{\mathrm{bn}}}{\varepsilon}, \\ \mathrm{Var}\left(C_t \mid \Omega\right) &\sim \frac{b\ \lambda}{r(\varepsilon+r)}\, e^{rt}\left(\frac{b\ \lambda}{r(\varepsilon+r)}\, e^{rt}+1\right) + \frac{B_0\ \lambda_{\mathrm{bn}}}{\varepsilon} \end{aligned} \tag{14}$$

for large $t$ - small $\lambda$.

***Exact results***

The asymptotic results discussed above perform extremely well for realistic parameter values and perfectly match the exact simulation results (see Figure S3). In principle, however, our mathematical framework can be used for other biomarkers - or entirely different applications - where the parameter values may not fall in the asymptotic regimes considered before. For this reason, we derive the exact distributions of the processes $A_t, B_t$ and $C_t$, for any given time $t$ and combinations of non-zero shedding rates.

<u>Probability generating functions</u>

In order to compute the exact distributions of $A_t$, $B_t$ and $C_t$, we first derive the joint probability generating functions of the processes $(A_t, C_t^A)$ and $(B_t, C_t^B)$. The marginal generating functions then follow directly, and provide the probability distributions of the individual processes by simple analytical or numerical inversion.

*Process* $(A_t, C_t^A)$

Our derivation for the process $(A_t, C_t^A)$ is based on previous results for a two-type branching process presented in refs. (*68, 69*). These studies assumed that cells of the second type also have the ability to reproduce, but shedding (or mutations) can happen only at wild-type cell death.

We consider instead the two-type branching process $(A_t, C_t^A)$ fully defined by the set of transitions in equation (8). For any given initial condition, we denote $P^*_{m,n}(t) = P((A_t, C_t^A) = (m, n) \mid (A_0, C_0^A) = *)$. The corresponding probability generating function is then defined as

$$\mathcal{P}^*(x,y,t) = \sum_{m,n\geq 0} x^m y^n P^*_{m,n}(t) \ .$$

The backward Kolmogorov equations for this process read

$$\begin{aligned} \frac{dP^{(1,0)}_{m,n}}{dt} &= (b-\lambda_2)P^{(2,0)}_{m,n} + (d-\lambda_0)\delta_{m,0}\delta_{n,0} \\ &\quad + \lambda_0 P^{(0,1)}_{m,n} + \lambda_1 P^{(1,1)}_{m,n} + \lambda_2 P^{(2,1)}_{m,n} - (b+d+\lambda_1)P^{(1,0)}_{m,n} \\ \frac{dP^{(0,1)}_{m,n}}{dt} &= \varepsilon\,\delta_{m,0}\delta_{n,0} - \varepsilon P^{(0,1)}_{m,n} \ . \end{aligned} \tag{15}$$

Hence, multiplying both sides of equation (15) by $x^m y^n$ and summing over all non-negative $m, n$ we find

$$\begin{aligned} \partial_t \mathcal{P}^{(1,0)} &= (b-\lambda_2)\mathcal{P}^{(2,0)} + (d-\lambda_0) \\ &\quad + \lambda_0 \mathcal{P}^{(0,1)} + \lambda_1 \mathcal{P}^{(1,1)} + \lambda_2 \mathcal{P}^{(2,1)} - (b+d+\lambda_1)\mathcal{P}^{(1,0)} \\ \partial_t \mathcal{P}^{(0,1)} &= \varepsilon - \varepsilon \mathcal{P}^{(0,1)} \ . \end{aligned}$$

Next, we observe that

$$\mathcal{P}^{(2,0)}(x,y,t) = \left[\mathcal{P}^{(1,0)}(x,y,t)\right]^2$$

because of the independence of the progenies of the two initial cells. By applying this property and introducing the notation

$$\mathcal{A}(x,y,t) = \mathcal{P}^{(1,0)}(x,y,t), \quad \mathcal{C}(x,y,t) = \mathcal{P}^{(0,1)}(x,y,t)$$

we reduce to the following system of first order differential equations

$$\partial_t \mathcal{A} = (b-\lambda_2)\mathcal{A}^2 + (d-\lambda_0) + \lambda_0\mathcal{C} + \lambda_1\mathcal{A}\mathcal{C} + \lambda_2\mathcal{A}^2\mathcal{C} - (b+d+\lambda_1)\mathcal{A} \tag{16}$$

$$\partial_t \mathcal{C} = \varepsilon - \varepsilon\mathcal{C} \tag{17}$$

with initial conditions

$$\mathcal{A}(x,y,0) = x \tag{18}$$

$$\mathcal{C}(x,y,0) = y \ . \tag{19}$$

Equation (17), subject to the initial condition (19), has solution

$$\mathcal{C}(x,y,t) = 1 + (y-1)e^{-\varepsilon t} \ . \tag{20}$$

Plugging this back into equation (16), we get

$$\partial_t \mathcal{A} = \left[b + \lambda_2(y-1)e^{-\varepsilon t}\right] \mathcal{A}^2 + \left[\lambda_1(y-1)e^{-\varepsilon t} - b - d\right] \mathcal{A} + \left[d + \lambda_0(y-1)e^{-\varepsilon t}\right] \ . \quad (21)$$

We apply the change of variables $s = e^{-\varepsilon t}$ and find

$$\partial_s \mathcal{A} = -\frac{[b + \lambda_2(y-1)s]}{\varepsilon s}\mathcal{A}^2 - \frac{[\lambda_1(y-1)s - b - d]}{\varepsilon s}\mathcal{A} - \frac{[d + \lambda_0(y-1)s]}{\varepsilon s} \ .$$

To ease the notation, we rewrite the equation above as

$$\partial_s \mathcal{A} = \frac{\alpha_2 + \beta_2 s}{s}\mathcal{A}^2 + \frac{\alpha_1 + \beta_1 s}{s}\mathcal{A} + \frac{\alpha_0 + \beta_0 s}{s} \quad (22)$$

where

$$\alpha_2 = -\frac{b}{\varepsilon}, \quad \alpha_1 = \frac{b+d}{\varepsilon}, \quad \alpha_0 = -\frac{d}{\varepsilon}$$
$$\beta_2 = -\frac{\lambda_2(y-1)}{\varepsilon}, \quad \beta_1 = -\frac{\lambda_1(y-1)}{\varepsilon}, \quad \beta_0 = -\frac{\lambda_0(y-1)}{\varepsilon} \ .$$

We are interested in the probability generating function $\mathcal{A}(x, y, t)$ for $t \geq 0$, that corresponds to $0 < s \leq 1$. Equation (22) is a Riccati equation, which can be reduced to a second order ODE. To do so, we define

$$X(x, y, s) = \frac{\alpha_2 + \beta_2 s}{s}\mathcal{A}(x, y, s)$$

which yields

$$\partial_s X = X^2 + \frac{(\alpha_2 + \beta_2 s)(\alpha_1 + \beta_1 s) - \alpha_2}{s(\alpha_2 + \beta_2 s)}X + \frac{(\alpha_2 + \beta_2 s)(\alpha_0 + \beta_0 s)}{s^2} \ . \quad (23)$$

Next, we set

$$X \equiv -\frac{\partial_s Y}{Y} \quad \Longrightarrow \quad \partial_s X = \frac{(\partial_s Y)^2}{Y^2} - \frac{\partial_s^2 Y}{Y}$$

so that equation (23) transforms into

$$\partial_s^2 Y - \frac{(\alpha_2 + \beta_2 s)(\alpha_1 + \beta_1 s) - \alpha_2}{s(\alpha_2 + \beta_2 s)}\partial_s Y + \frac{(\alpha_2 + \beta_2 s)(\alpha_0 + \beta_0 s)}{s^2}Y = 0 \ . \quad (24)$$

Equation (24) is a second order linear differential equation with rational coefficients. It has three singular points: $s = 0$ and $s = -\frac{\alpha_2}{\beta_2}$ being regular and $s = \infty$ being irregular with rank 1. Hence, equation (24) is a single confluent Heun equation (*70*). To bring it into standard form, we first move the non-zero regular singularity to 1 through the change of variable $z = -\frac{\beta_2}{\alpha_2} s$. This leads to

$$\partial_z^2 Y + \frac{(z-1)\left(\frac{\alpha_2\beta_1}{\beta_2} z - \alpha_1\right) - 1}{z(z-1)} \partial_z Y + \frac{\alpha_2 (z-1)\left(\frac{\alpha_2\beta_0}{\beta_2} z - \alpha_0\right)}{z^2} Y = 0 \ . \tag{25}$$

We now look for a solution to equation (25) of the form

$$Y(z) = z^m e^{nz} f(z) \tag{26}$$

which implies

$$\begin{aligned} \partial_z Y &= z^{m-1} e^{nz} \left[(m+nz) f + z f'\right] \\ \partial_z^2 Y &= z^{m-2} e^{nz} \left\{\left[(m+nz)^2 - m\right] f + 2z(m+nz) f' + z^2 f''\right\} \ . \end{aligned}$$

Substituting these expressions into equation (25) and rearranging, we obtain the following differential equation for $f(z)$

$$f'' + \frac{P(z)}{z(z-1)} f' + \frac{Q(z)}{z^2(z-1)} f = 0 \tag{27}$$

where $P(z)$ and $Q(z)$ are two polynomials of second and third degree in $z$, respectively. However, it is easily checked that by taking

$$m = -\alpha_2, \quad n = -\frac{\alpha_2}{2\beta_2}\left[\beta_1 + \sqrt{\beta_1^2 - 4\beta_0\beta_2}\right] \tag{28}$$

the first and last coefficients of the polynomial $Q(z)$ become zero. Hence, for these values equation (27) can be written as

$$f'' + \left(\frac{\gamma}{z} + \frac{\delta}{z-1} + \eta\right) f' + \frac{\omega z + \rho}{z(z-1)} f = 0 \ . \tag{29}$$

Here, the parameters $\gamma, \delta$ and $\eta$ follow from the decomposition in partial fractions of $\frac{P(z)}{z(z-1)}$ while $\omega$ and $\rho$ correspond to the coefficients of second and first degree terms in $Q(z)$, respectively - when $m$ and $n$ are set as in equation (28). Explicit expressions for all these parameters in terms of the original rates are given by

$$\begin{aligned}
m &= \frac{b}{\varepsilon}, \qquad n = \frac{b}{2\lambda_2\varepsilon}\left(\lambda_1 - \sqrt{\lambda_1^2 - 4\lambda_0\lambda_2}\right) \\
\gamma &= \frac{b-d}{\varepsilon} + 1\,, \qquad \delta = -1\,, \qquad \eta = \frac{b}{\lambda_2\varepsilon}\sqrt{\lambda_1^2 - 4\lambda_0\lambda_2} \\
\omega &= -\frac{b}{2\lambda_2\varepsilon^2}\left[(b+d)\lambda_1 - (b-d)\sqrt{\lambda_1^2 - 4\lambda_0\lambda_2} + 2(b\lambda_0 + d\lambda_2)\right] \\
\rho &= \frac{b}{2\lambda_2\varepsilon^2}\left\{(b+d-\varepsilon)\lambda_1 - (b-d+\varepsilon)\sqrt{\lambda_1^2 - 4\lambda_0\lambda_2} + 2[b\lambda_0 + (d-\varepsilon)\lambda_2]\right\}\ .
\end{aligned} \tag{30}$$

Now, equation (29) is the standard form of the single confluent Heun equation. Solutions to such a second order differential equation are called confluent Heun functions and depend on six arguments - the five parameters of the equation and the independent variable $z$. A standardized package for numerical and symbolical computations involving Heun functions is currently provided only by Maple (*71*), which includes in particular the procedures `HeunC` and `HeunCPrime` for the evaluation of confluent Heun functions and their $z$ derivative, respectively. Constistently with these implementations the general solution of equation (29) can be written as

$$f(z) = h_1(z) + Dz^{1-\gamma}h_2(z) \tag{31}$$

where

$$\begin{aligned}
h_1(z) &= \texttt{HeunC}\left(\eta, 1-\gamma, \delta-1, \omega - \frac{\eta(\gamma+\delta)}{2}, \rho + \frac{1-\gamma(\delta-\eta)}{2}, z\right) \\
h_2(z) &= \texttt{HeunC}\left(\eta, \gamma-1, \delta-1, \omega - \frac{\eta(\gamma+\delta)}{2}, \rho + \frac{1-\gamma(\delta-\eta)}{2}, z\right)
\end{aligned}$$

are the confluent Heun functions uniquely determined by the following initial conditions

$$\begin{aligned}
h_1(0) &= 1, \qquad h_1'(0) = \frac{\rho}{\gamma} \\
h_2(0) &= 1, \qquad h_2'(0) = \frac{(\gamma-1)(\delta-\eta)-\rho}{\gamma-2}\ .
\end{aligned} \tag{32}$$

Here $h_i'$ denotes the $z$ derivative of the confluent Heun functions $h_i$. As mentioned before these functions are implemented by the Maple procedure `HeunCPrime` and uniquely determined by an additional condition on $h_i''(0)$, too cumbersome to be reported here. Also notice that the expression in equation (31) contains only one integrating constant, $D$, as our derivation spawns from a first order differential equation. By plugging such expression back into equation (26), the solution to equation (25) becomes

$$Y(z) = z^m e^{nz} f(z) = z^m e^{nz} h_1(z) + D z^{1+m-\gamma} e^{nz} h_2(z) \ .$$

Next, we denote the derivative of $f(z)$ as

$$g(z) = f'(z) = h_1'(z) + D z^{-\gamma}[(1-\gamma)h_2(z) + z h_2'(z)] \ . \tag{33}$$

Recalling that $z = -\frac{\beta_2}{\alpha_2} s = ks$ and that $X(x,y,s) = -\frac{\partial_s Y}{Y}$, we find

$$X(x,y,s) = -\frac{(m+nks)f(ks) + ksg(ks)}{sf(ks)} \ . \tag{34}$$

Notice that in terms of the original parameters of our model the coefficient $k$ is given by $k = q_b(y-1)$ and thus depends on $y$. We can now apply the initial condition for $X(x,y,s)$ to find the value of the constant $D$. Since $s = e^{-\varepsilon t}$ and $X(x,y,s) = \frac{\alpha_2+\beta_2 s}{s}\mathcal{A}(x,y,s)$, the initial condition $\mathcal{A}(x,y,t=0) = x$ translates to $X(x,y,s=1) = (\alpha_2+\beta_2)x = \alpha_2(1-k)x$. Therefore, equation (34) at $s=1$ implies

$$\alpha_2(1-k)x = -\frac{(m+nk)f(k) + kg(k)}{f(k)} \ .$$

Substituting the expressions for $f$ and its derivative (equations (31) and (33), respectively) and solving for $D$, we find

$$D = -\frac{(m+nk+\alpha_2 x - k\alpha_2 x)h_1(k) + kh_1'(k)}{k^{1-\gamma}[(m+nk+\alpha_2 x - k\alpha_2 x + 1 - \gamma)h_2(k) + kh_2'(k)]} \ .$$

Plugging this value back into equation (34), we find an expression for $X$ in terms of the functions $h_i(s)$ and $h_i'(s)$. Multiplying this expression by $\frac{s}{\alpha_2(1-ks)}$ and substituting $s = e^{-\varepsilon t}$, we obtain

$$\mathcal{A}(x,y,t) = \frac{e^{rt}K_1(x,y)\phi_2(y,t) - K_2(x,y)\phi_1(y,t)}{e^{rt}K_1(x,y)\psi_1(y,t) - K_2(x,y)\psi_2(y,t)} \tag{35}$$

where

$$
\begin{aligned}
K_1(x,y) &= \left\{\frac{b - x[b+\lambda_2(y-1)]}{\varepsilon} + nk\right\} h_1(k) + kh_1'(k) \\
K_2(x,y) &= \left\{\frac{d - x[b+\lambda_2(y-1)]}{\varepsilon} + nk\right\} h_2(k) + kh_2'(k) \\
\phi_1(y,t) &= \left[\frac{bnke^{-\varepsilon t}}{\varepsilon} + 1\right] h_1(ke^{-\varepsilon t}) + \frac{\varepsilon k}{b} e^{-\varepsilon t} h_1'(ke^{-\varepsilon t}) \\
\phi_2(y,t) &= \left[\frac{bnke^{-\varepsilon t}}{\varepsilon} + \frac{d}{b}\right] h_2(ke^{-\varepsilon t}) + \frac{\varepsilon k}{b} e^{-\varepsilon t} h_2'(ke^{-\varepsilon t}) \\
\psi_1(y,t) &= (1 - ke^{-\varepsilon t}) h_1(ke^{-\varepsilon t}) \\
\psi_2(y,t) &= (1 - ke^{-\varepsilon t}) h_2(ke^{-\varepsilon t}) \ .
\end{aligned} \tag{36}
$$

This is the joint probability generating function of the processes $A_t$ and $C_t^A$, starting from one $A$ cell at time $t = 0$.

To find the marginal generating function

$$
\mathscr{A}(x,t) = \sum_{m=0}^{\infty} \mathrm{P}(A_t = m \mid (A_0, C_0^A) = (1,0)) x^m
$$

we take the limit for $y \to 1$ in equation (35). Given the conditions in equation (32), this yields

$$
\mathscr{A}(x,t) = \frac{e^{-rt}(x-\delta) + \delta(1-x)}{e^{-rt}(x-\delta) + (1-x)} \ . \tag{37}
$$

As expected, this coincides with the probability generating function of a supercritical birth death process with net growth rate $r = b - d > 0$ and extinction probability $\delta = d/b < 1$. The exact distribution for the process $A_t$ then follows by inverting $\mathscr{A}(x,t)$ and is given by

$$
\mathrm{P}(A_t = m) = \frac{1}{m!} \frac{\partial^m \mathscr{A}(x,t)}{\partial x^m} \Big|_{x=0} = \begin{cases} \frac{\delta}{S(t)} & \text{if } m = 0 \\ \left(1 - \frac{\delta}{S(t)}\right)(S(t)-1)S(t)^{-m} & \text{if } m \geq 1 \end{cases} \tag{38}
$$

where $S(t) = \frac{1-\delta e^{-rt}}{1-e^{-rt}}$. Similarly, the generating function

$$
\mathscr{C}^A(y,t) = \sum_{n=0}^{\infty} \mathrm{P}(C_t^A = n \mid (A_0, C_0^A) = (1,0)) y^n
$$

is derived by taking the limit for $x \to 1$ in equation (35). The expression for $\mathscr{C}^A$

slightly simplifies to

$$\mathscr{C}^A(y,t) = \frac{e^{rt}\bar{K}_1(y)\phi_2(y,t) - \bar{K}_2(y)\phi_1(y,t)}{e^{rt}\bar{K}_1(y)\psi_1(y,t) - \bar{K}_2(y)\psi_2(y,t)} \tag{39}$$

where

$$\bar{K}_1(y) = \left\{nk - \frac{\lambda_2(y-1)}{\varepsilon}\right\} h_1(k) + kh_1'(k)$$
$$\bar{K}_2(y) = \left\{nk - \frac{b-d+\lambda_2(y-1)}{\varepsilon}\right\} h_2(k) + kh_2'(k)$$

and the functions $\phi_1, \phi_2, \psi_1, \psi_2$ are defined as in equation (36). The function $\mathscr{C}^A(y,t)$ can be inverted numerically (see discussion below on inversion techniques) to find the distribution of the process $C_t^A$.

We present the particular form that these results assume when one or more of the parameters $q_d, \lambda_1, q_b$ are zero. In general, among the nonzero shedding rates, the one with the highest index determines the special functions involved in the generating function $\mathcal{A}(x,y,t)$. As shown above, these are confluent Heun functions when $\lambda_2 > 0$, while they become confluent hypergeometric functions when $\lambda_2 = 0$ and $\lambda_1 > 0$ and Bessel functions of the first kind when $\lambda_2 = 0, \lambda_1 = 0$ and $\lambda_0 > 0$. The remaining cases follow by straightforward substitution from these three. We provide a sketch of how the previous derivation can be adapted to the cases $\lambda_2 = 0, \lambda_1 > 0$ and $\lambda_2 = 0, \lambda_1 = 0, \lambda_0 > 0$. The steps up until equation (24) are valid for all scenarios, which will be the starting point of the adapted derivations. Also notice that for all these cases the marginal probability generating function $\mathscr{A}(y,t)$ remains unchanged, as the evolution of the process $A_t$ is not affected by its shedding activity.

$q_b = 0, \lambda_1 > 0$

When $q_b = 0$, we have $\beta_2 = 0$. Hence, equation (24) becomes

$$\partial_s^2 Y - \frac{\beta_1 s + \alpha_1 - 1}{s}\partial_s Y + \frac{\alpha_2(\alpha_0 + \beta_0 s)}{s^2} Y = 0$$

By seeking directly a solution of the form $Y(s) = s^{-\alpha_2} f(s)$ and then applying the change of variables $z = \beta_1 s$, the equation above reduces to the standard form of Kummer's equation (*72*)

$$zf''(z) + (\gamma - z)f'(z) - \omega f(z) = 0 \tag{40}$$

where

$$\gamma = \alpha_0 - \alpha_2 + 1$$
$$c = \beta_1$$
$$\omega = -\alpha_2 \left( \frac{\beta_0}{\beta_1} + 1 \right) \ .$$

Including again only one integrating constant $D$, its solution can be expressed in terms of the confluent hypergeometric function ${}_1F_1$ as

$$f(z) = {}_1F_1(\omega, \gamma, z) + Dz^{1-\gamma}{}_1F_1(\omega - \gamma + 1, 2 - \gamma, z)$$

Substituting back we immediately find an expression for the solution $Y(s)$ in terms of

$$h_1(s) = {}_1F_1(\omega, \gamma, cs), \quad h_2(s) = {}_1F_1(\omega - \gamma + 1, 2 - \gamma, cs) \ .$$

Furthermore, since

$$\frac{\partial}{\partial z}{}_1F_1(a, b, z) = \frac{a}{b}{}_1F_1(a + 1, b + 1, z)$$

the derivative of $Y(s)$ can be computed in terms of the functions $h_1, h_2, g_1$ and $g_2$, where

$$g_1(s) = {}_1F_1(\omega + 1, \gamma + 1, cs), \quad g_2(s) = {}_1F_1(\omega - \gamma + 2, 3 - \gamma, cs) \ .$$

Joining these results together, we find an expression for $X(s)$, in terms of the same functions. This expression still depends on the integrating constant $D$, which is determined by the initial condition $X(x, y, s = 1) = \alpha_2 x$. Then, multiplying $X(x, y, s)$ by $\frac{s}{\alpha_2}$ and substituting $s = e^{-\varepsilon t}$ we eventually find

$$\mathcal{A}(x, y, t) = \frac{e^{rt}K_1(x, y)\phi_2(y, t) - K_2(x, y)\phi_1(y, t)}{e^{rt}K_1(x, y)\psi_1(y, t) - K_2(x, y)\psi_2(y, t)} \tag{41}$$

where

$$\begin{aligned}
K_1(x,y) &= (b-d+e)(1-x)h_1(y,0) - (\lambda_1+\lambda_0)(y-1)g_1(y,0)\\
K_2(x,y) &= (b-d-e)(d-bx)h_2(y,0) + (d\lambda_1+b\lambda_0)(y-1)g_2(y,0)\\
\phi_1(y,t) &= (b-d+e)h_1(y,t) - (\lambda_1+\lambda_0)(y-1)e^{-\varepsilon t}g_1(y,t)\\
\phi_2(y,t) &= d(b-d-e)h_2(y,t) + (d\lambda_1+b\lambda_0)(y-1)e^{-\varepsilon t}g_2(y,t)\\
\psi_1(y,t) &= (b-d+e)h_1(y,t)\\
\psi_2(y,t) &= b(b-d-e)h_2(y,t)\ .
\end{aligned}$$

The marginal probability generating function for the process $C_t^A$ becomes

$$\mathscr{C}^A(y,t) = \frac{e^{rt}\bar{K}_1(y)\phi_2(y,t) - \bar{K}_2(y)\phi_1(y,t)}{e^{rt}\bar{K}_1(y)\psi_1(y,t) - \bar{K}_2(y)\psi_2(y,t)} \tag{42}$$

where

$$\begin{aligned}
\bar{K}_1(y) &= -(\lambda_1+\lambda_0)(y-1)g_1(y,0)\\
\bar{K}_2(y) &= (b-d-e)(d-b)h_2(y,0) + (d\lambda_1+b\lambda_0)(y-1)g_2(y,0)\ .
\end{aligned}$$

$q_b = 0, \lambda_1 = 0$

If we additionally have $\lambda_1 = 0$, equation (24) becomes

$$\partial_s^2 Y - \frac{\alpha_1 - 1}{s}\partial_s Y + \frac{\alpha_2(\alpha_0+\beta_0 s)}{s^2}Y = 0\ .$$

Through the change of variables $z = 2\sqrt{\alpha_2\beta_0 s} = c\sqrt{s}$ and by seeking a solution of the form $Y(z) = z^{\alpha_1}f(z)$, we reduce to the Bessel equation (*72*)

$$z^2 f'' + zf' + [z^2 - (\alpha_2-\alpha_0)^2]f = 0\ . \tag{43}$$

The solution to equation (43) can be expressed as a Bessel function of the first kind $J_\nu(z)$ as

$$f(z) = J_{\alpha_2-\alpha_0}(z) + DJ_{\alpha_0-\alpha_2}(z)\ .$$

This allows us to write $Y(s)$ in terms of $h_1(s) = J_{\alpha_0-\alpha_2}(c\sqrt{s})$ and $h_2(s) = J_{\alpha_2-\alpha_0}(c\sqrt{s})$ and since

$$\partial_z J_w(z) = \frac{w}{z}J_w(z) - J_{w+1}(z)\ ,$$

we can compute $Y(s)$ derivative in terms of the functions $h_1, h_2, g_1$ and $g_2$, where

$$g_1(s) = J_{\alpha_0-\alpha_2+1}(c\sqrt{s}), \quad g_2(s) = J_{\alpha_2-\alpha_0+1}(c\sqrt{s}) \ .$$

By combining these expressions, we first derive $X(s)$, and then find the integrating constant $D$ by applying the initial condition $X(x, y, s = 1) = \alpha_2 x$. Multiplying by $\frac{s}{\alpha_2}$ and substituting $s = e^{-\varepsilon t}$, we find

$$\mathcal{A}(x, y, t) = \frac{K_1(x, y)\phi_2(y, t) - K_2(x, y)\phi_1(y, t)}{K_1(x, y)\psi_2(y, t) - K_2(x, y)\psi_1(y, t)} \tag{44}$$

where

$$\begin{aligned}
K_1(x, y) &= (bx - b)h_1(y, 0) + \sqrt{b\lambda_0(y-1)}g_1(y, 0)\\
K_2(x, y) &= (bx - d)h_2(y, 0) + \sqrt{b\lambda_0(y-1)}g_2(y, 0)\\
\phi_1(y, t) &= bh_1(y, t) - \sqrt{b\lambda_0(y-1)}e^{-\frac{\varepsilon}{2}t}g_1(y, t)\\
\phi_2(y, t) &= dh_2(y, t) - \sqrt{b\lambda_0(y-1)}e^{-\frac{\varepsilon}{2}t}g_2(y, t)\\
\psi_1(y, t) &= bh_1(t)\\
\psi_2(y, t) &= bh_2(t) \ .
\end{aligned}$$

From here we can recover the usual marginal probability generating function for the process $A_t$ by considering the expansions of Bessel functions $J_w(z)$ around $z = 0$. The marginal probability generating function for the process $C_t^A$ is instead given by

$$\mathscr{C}^A(y, t) = \frac{\bar{K}_1(y)\phi_2(y, t) - \bar{K}_2(y)\phi_1(y, t)}{\bar{K}_1(y)\psi_1(y, t) - \bar{K}_2(y)\psi_2(y, t)} \tag{45}$$

where

$$\begin{aligned}
\bar{K}_1(y) &= \sqrt{b\lambda_0(y-1)}g_1(y, 0)\\
\bar{K}_2(y) &= (b - d)h_2(y, 0) + \sqrt{b\lambda_0(y-1)}g_2(y, 0) \ .
\end{aligned}$$

Finally, we remark that in all the derivations above, there are special cases of the equation $f(z)$ that require extra attention. Namely, if some of the equation coefficients are integers, then its general solution assumes a slightly different expression than the one reported. However, it is unlikely that real estimates lead to such special cases, which we therefore do not discuss in detail here.

*Process* $(B_t, C_t^B)$

We derive exact results for the size distributions of the processes $B_t$ and $C_t^B$. So far, we assumed a constant population of benign cells and studied the asymptotic behaviour of the biomarker amount they shed. Here, we first present additional details about this case, providing the exact generating function of $C_t^B$ at a given time $t$. Later, we show how a derivation similar to that employed for the process $(A_t, C_t^A)$ can be used when $B_t$ is modeled by a critical branching birth-death process.

Constant growth

In the derivation of asymptotic results for our model, we already mentioned that if $B_t$ is constant then $C_t^B$ is a branching pure-death process with immigration and its exact probability generating function is given by equation (3). In our setup, we also assume that $C_t^B$ is originally at equilibrium, i.e. that it starts at the expected value of its large time asymptotic distribution. Since we observed that in this limit $C_t^B$ converges to a Poisson random variable with mean $\frac{\lambda_{\text{bn}}}{\varepsilon}B_0$, we set $C_0^B = \frac{\lambda_{\text{bn}}}{\varepsilon}B_0$. With this initial condition, equation (3) becomes

$$\mathscr{C}^B(y,t) = \left\{\left[1 + (y-1)e^{-\varepsilon t}\right] e^{(y-1)(1-e^{-\varepsilon t})}\right\}^{\frac{\lambda_{\text{bn}}}{\varepsilon}B_0} .$$

Numerical inversion of this function then provides the exact distribution of the process $C_t^B$ at any given time $t$.

Critical growth

In our model $B_t$ denotes the number of benign cells at time $t$ with the ability to shed the biomarker in the bloodstream. So far we have assumed that this number stays constant over time, but more generally this number can fluctuate around a constant average value. For this reason, $B_t$ can be modeled as a critical branching birth death process, i.e. as a process defined like $A_t$ but with the same birth and death rates $b_{\text{bn}} = d_{\text{bn}}$. Under this assumption, and assuming that the shedding probabilities at cell apoptosis and proliferation are the same for benign and malignant cells, the set of transitions characterizing the two-type process $(B_t, C_t^B)$ are

$$\begin{array}{lll} B \longrightarrow BBC & \text{rate } b_{\text{bn}}\, q_b & \text{rate } \lambda_{\text{bn},2} \\ B \longrightarrow BB & \text{rate } b_{\text{bn}}\, (1-q_b) & \text{rate } b_{\text{bn}} - \lambda_{\text{bn},2} \\ B \longrightarrow BC & \text{rate } \lambda_{\text{bn},1} & \text{rate } \lambda_{\text{bn},1} \\ B \longrightarrow C & \text{rate } b_{\text{bn}}\, q_d & \text{rate } \lambda_{\text{bn},0} \\ B \longrightarrow \emptyset & \text{rate } b_{\text{bn}}\, (1-q_d) & \text{rate } b_{\text{bn}} - \lambda_{\text{bn},1} \\ C \longrightarrow \emptyset & \text{rate } \varepsilon & \text{rate } \varepsilon\ . \end{array}$$

The exact probability generating functions for this process can be computed through a derivation similar to that shown for $(A_t, C_t^A)$. Briefly, we can redefine $\mathcal{P}^*$ as

$$\mathcal{P}^*(x,y,t) = \sum_{m,n\geq 0} x^m y^n \, \mathrm{P}\left(\left(B_t, C_t^B\right) = (m,n) \mid \left(B_0, C_0^B\right) = *\right)$$

and set

$$\mathcal{B}(x,y,t) = \mathcal{P}^{(1,0)}(x,y,t), \qquad \mathcal{C}'(x,y,t) = \mathcal{P}^{(0,1)}(x,y,t)\ .$$

As biomarker units are eliminated from the bloodstream at the same rate regardless of the cell type that shed them, we have $\mathcal{C}'(x,y,t) = \mathcal{C}(x,y,t)$, where $\mathcal{C}(x,y,t)$ is given by equation (20). However, the function $\mathcal{B}(x,y,t)$ does not follow straightforwardly from $\mathcal{A}(x,y,t)$ by simply substituting the dashed rates and taking the limit as $d_{\text{bn}} \to b_{\text{bn}}$. The reason is that in this limit the two linearly independent solutions $h_1(z)$ and $h_2(z)$ of the reduced equation become the same and so a different set of solutions has to be chosen. Once these are found, the subsequent steps can be repeated and an explicit expression for $\mathcal{B}(x,y,t)$ can be derived. The exact probability generating function for the process $(B_t, C_t^B)$ is given by

$$\mathcal{P}^{(B_0,C_0^B)}(x,y,t) = \mathcal{B}(x,y,t)^{B_0}\mathcal{C}(x,y,t)^{C_0^B}\ .$$

The marginal generating functions for the two single processes follow from the $y \to 1$ and $x \to 1$ limits, respectively. In particular, for any combination of non-zero shedding rates the former is equal to

$$\mathscr{B}(x,t) = \left(1 + \frac{x-1}{1 - d_{\text{bn}}t(x-1)}\right)^{B_0}$$

which coincides with the probability generating function of a critical branching birth death process with death rate $d_{\text{bn}}$ and $B_0$ initial individuals. This function is analytically invertible, even though the terms of the probability mass function of $B_t$

can only be expressed as finite sums

$$\mathrm{P}(B_t = m \mid B_0) = \frac{B_0 \left(\frac{d_{\mathrm{bn}} t}{d_{\mathrm{bn}} t+1}\right)^{B_0}}{m(d_{\mathrm{bn}} t)^m (1+d_{\mathrm{bn}} t)^m} \sum_{j=0}^{m-1} \binom{B_0 - 1}{B_0 - m + j} \binom{m}{j} (d_{\mathrm{bn}} t)^{2j} \ .$$

The marginal generating function of $C_t^B$ can instead be inverted numerically to derive the exact probability distribution of the process. However, while the above described computations are feasible, modeling $B_t$ as a critical branching birth death process leads to tedious complications and does not practically change the dynamics of the model. The complications are related to the fact that a critical branching process eventually gets extinct with probability one. We are generally interested in the large time limit distribution of $B_t$, but this limit converges to a point mass at zero and conditioning on $B_t$ eventual survival does not make sense either. Moreover, since $B_t$ starts with a very large number of cells, the time required by $B_t$ to become extinct would be unrealistically long. Indeed, the expected time to extinction for a critical branching process is infinite, and using the expression above with $d_{\mathrm{bn}} = 0.1$, we find that $\mathrm{P}(B_{10000 \text{ yrs}} = 0 \mid B_0 = 10^7) \approx 10^{-12}$. For the same principle, the probability that $B_t$ exhibits significant deviations from $B_0$ within human lifetime is very small. To quantify the probability of such fluctuations, we exploit Chebyshev inequality: using again $d_{\mathrm{bn}} = 0.1$ and $B_0 = 10^7$, we find that $\mathrm{P}(|B_{55 \text{ yrs}} - 10^7| \geq 2 \times 10^6) \leq 0.01$, which says that even after waiting 55 years, the probability of observing a deviation of at least 20% from the original population size would still be lower than 1%.

*Process $C_t$ and numerical inversion*

Given the independence of biomarker shedding from benign and malignant cells, the probability generating function of the process $C_t = C_t^A + C_t^B$ is equal to

$$\mathscr{C}(y,t) = \mathscr{C}^A(y,t)\mathscr{C}^B(y,t)$$

where the two functions on the right hand side are given by equation (39) (see also equations (42) and (45) for special cases) and equation (3) (assuming $B_t$ is constant), respectively. As for many other functions derived before, $\mathscr{C}(y,t)$ can be numerically inverted to find the exact distribution of the process $C_t$.

We briefly discuss some inversion techniques. The goal of the inversion procedure is to derive the probability mass function of a stochastic process $\chi_t$ from its generating function $G(z,t) = \sum_{k=0}^{\infty} \mathrm{P}(\chi_t = k) z^k$ as

$$\mathrm{P}(\chi_t = k) = \frac{1}{k!} \frac{\partial^k G(z,t)}{\partial z^k}\Big|_{z=0} \, . \tag{46}$$

In a few cases, the $k$-th derivative of the generating function is explicitly computable, thus allowing us to find an analytic formula for the probability mass function of $\chi_t$. The expression for $\mathscr{A}(x,t)$ given by equation (37) is an example of such a generating function and from it we directly derived the probabilities $\mathrm{P}(A_t = m)$ (see equation (38)). For most generating functions, however, this analytical inversion is not feasible and we have to compute the probability mass function numerically. Algorithms for this numerical procedure are based on techniques for series expansions of analytical functions, which are implemented as `Series` in both Mathematica (see also `NSeries`) and Maple. For more details on the numerical inversion of probability generating functions, we refer to refs. (*73, 74*).

Conditional distributions

In our model summary we pointed out that we are interested in the probability distributions of the processes $A_t, B_t$ and $C_t$ conditional on $A_t$ non-extinction. Therefore, the asymptotic results derived before are conditioned on the event of $A_t$ eventual survival, which we denoted as $\Omega$. The same results were also expressed in terms of the primary tumor size, by computing the asymptotic distributions of the processes involved at the time when $A_t = m$, for $m$ large. Here we show how the same kind of conditioning can be applied to the exact probability mass functions derived above.

We note that all the probabilities involved in the following derivations are conditioned on the usual initial values of the processes $A_t, B_t$ and $C_t$. For clarity, we do not introduce a new notation for such conditioning, and hereafter we implicitly denote

$$\mathrm{P}\,(\,\cdot\,) = \mathrm{P}\left(\,\cdot\,\mid \left(A_0, B_0, C_0^A, C_0^B\right) = \left(1, B_0, 0, \frac{\lambda_{\mathrm{bn}}}{\varepsilon} B_0\right)\right) \, .$$

*Conditioning on $A_t$ survival*

As we are dealing with exact distributions at a given time, we condition on the event of $A_t$ survival up to time $t$, that is

$$\Omega_t = \{A_t > 0\} \, .$$

We denote $\delta_t = \mathrm{P}(\Omega_t^c) = \mathrm{P}(A_t = 0)$. This probability is equal to $\mathscr{A}(0,t)$, so we get

$$\delta_t = \frac{\delta - \delta e^{-rt}}{1 - \delta e^{-rt}} \ .$$

Bayes theorem yields

$$\mathrm{P}\left(A_t = m \mid \Omega_t\right) = \frac{\mathrm{P}\left(A_t = m\right)}{\mathrm{P}\left(\Omega_t\right)} \cdot \mathrm{P}\left(\Omega_t \mid A_t = m\right)$$

for any $m \geq 0$. The term $\mathrm{P}\left(\Omega_t \mid A_t = m\right)$, however, is equal to 0 for $m = 0$ and to 1 for every $m \geq 1$. Hence, we get

$$\mathrm{P}\left(A_t = m \mid \Omega_t\right) = \frac{\mathrm{P}\left(A_t = m\right)}{1 - \delta_t}$$

for every $m \geq 1$, and 0 otherwise.

Similar steps allow us to compute $\mathrm{P}\left(C_t^A = n \mid \Omega_t\right)$. In this case, we find

$$\mathrm{P}\left(C_t^A = n \mid \Omega_t\right) = \frac{\mathrm{P}\left(C_t^A = n, \Omega_t\right)}{\mathrm{P}\left(\Omega_t\right)} = \frac{\mathrm{P}\left(C_t^A = n\right) - \mathrm{P}\left(\left(A_t, C_t^A\right) = (0,n)\right)}{1 - \delta_t} \ . \tag{47}$$

The terms $\mathrm{P}(C_t^A = n)$ follow from the probability generating function $\mathscr{C}^A(y,t)$. Similarly, the probabilities $\mathrm{P}\left(\left(A_t, C_t^A\right) = (0,n)\right)$ are obtained by inverting the function $\mathcal{A}(0,y,t)$.

*Conditioning on one process size*

The distribution of the total biomarker amount present in the bloodstream at time $t$ conditioned on the primary tumor size at that time is

$$\begin{aligned}\mathrm{P}(C_t = n \mid A_t = m) &= \sum_{i=0}^{n} \mathrm{P}\left(C_t^B = i\right) \mathrm{P}\left(C_t^A = n - i \mid A_t = m\right) \\ &= \frac{\sum_{i=0}^{n} \mathrm{P}\left(C_t^B = i\right) \mathrm{P}\left(A_t = m, C_t^A = n - i\right)}{\mathrm{P}(A_t = m)} \ .\end{aligned}$$

The terms $\mathrm{P}\left(A_t = m, C_t^A = n - i\right)$ follow from inverting the joint probability generating function $\mathcal{A}(x,y,t)$. Moreover, in this case, if we consider a strictly positive primary tumor size $m$, conditioning on $A_t$ survival is not necessary as $\{A_t = m\} \subset \Omega_t$.

We note that once the exact probability mass functions are known one can conversely condition on the total biomarker amount present and ask how this affects the primary tumor size distributions. To see this, we write

$$
\begin{aligned}
\mathrm{P}(A_t = m \mid C_t = n) &= \frac{\mathrm{P}(A_t = m, C_t = n)}{\mathrm{P}(C_t = n)} \\
&= \frac{\sum_{i=0}^{n} \mathrm{P}\left(C_t^B = i\right) \mathrm{P}\left(A_t = m, C_t^A = n - i\right)}{\sum_{i=0}^{n} \mathrm{P}\left(C_t^B = i\right) \mathrm{P}\left(C_t^A = n - i\right)} \ .
\end{aligned}
$$

By further conditioning on $A_t$ survival up to $t$, we obtain

$$
\mathrm{P}(A_t = m \mid C_t = n, \Omega_t) = \frac{\mathrm{P}(A_t = m, \Omega_t \mid C_t = n)}{\mathrm{P}(\Omega_t \mid C_t = n)} = \frac{(1-\delta_t)\,\mathrm{P}(A_t = m \mid C_t = n)}{\mathrm{P}(C_t = n \mid \Omega_t)\,\mathrm{P}(C_t = n)} \ .
$$

Finally, we recall that because $(B_t, C_t^B)$ is independent of the primary tumor growth dynamics, any kind of conditioning on $A_t$ size has no effect on the $B_t$ and $C_t^B$ probability distributions.

<u>Expected values</u>

From equation (46) we recover the well known formulas for the mean and variance of a discrete state stochastic process $\chi_t$ with probability generating function $G(z,t)$

$$
\mathrm{E}[\chi_t] = \partial_z\, G(z,t)_{|z=1}, \quad \mathrm{Var}(\chi_t) = \partial_z^2\, G(z,t)_{|z=1} + \mathrm{E}[\chi_t] - \mathrm{E}[\chi_t]^2 \ .
$$

These properties allow us to derive analytical expression for the expected value and variance of the processes $A_t, B_t$ and $C_t$ at any given time $t$. In this section, we summarize these expressions, implicitly conditioning on $(A_0, C_0^A) = (1,0)$ and $C_0^B = \frac{\lambda_{\mathrm{bn}}}{\varepsilon} B_0$.

For the malignant cancer cell population, we have

$$
\mathrm{E}[A_t] = e^{rt}, \qquad \mathrm{Var}(A_t) = \frac{1+\delta}{1-\delta}\, e^{rt}(e^{rt} - 1)
$$

and

$$
\mathrm{E}[A_t \mid \Omega_t] = \frac{e^{rt} - \delta}{1-\delta}, \quad \mathrm{Var}(A_t \mid \Omega_t) = \frac{(1-\delta e^{-rt})(1+2\delta)e^{rt}(e^{rt}-1)}{(1-\delta)^2} \ .
$$

When the benign population is modeled by a critical branching birth death process, we find

$$\mathrm{E}[B_t] \equiv B_0, \qquad \mathrm{Var}(B_t) = 2d_{\mathrm{bn}} B_0 t \ .$$

In this case, conditioning on $A_t$ survival has no effect as $A_t$ and $B_t$ are independent processes.

As for the total amount of circulating biomarker, from the functions $\mathscr{C}^A(y,t)$ and $\mathscr{C}^B(y,t)$ we get

$$\mathrm{E}[C_t^A] = \frac{\lambda\left(e^{rt} - e^{-\varepsilon t}\right)}{r+\varepsilon}, \qquad \mathrm{E}[C_t^B] \equiv \frac{\lambda_{\mathrm{bn}}}{\varepsilon} B_0 \ .$$

The expression for the latter mean is the same for $B_t$ modeled as a constant population or as a critical branching birth death process. Furthermore, in our setup such expectation does not depend on $t$ as we start the process $C_t^B$ at its equilibrium. Combining the previous results, we find

$$\mathrm{E}[C_t] = \frac{\lambda\left(e^{rt} - e^{-\varepsilon t}\right)}{r+\varepsilon} + \frac{\lambda_{\mathrm{bn}}}{\varepsilon} B_0 \ .$$

The variance of $C_t$ and expressions for its first moments conditional on $A_t$ survival can be obtained from equation (47).

Sampling scheme

By combining the previous results, we can now derive the exact probability distribution for the total biomarker amount present in a blood sample of a given volume at time $t$. To this end, we first assume that at time $t$ there is a fixed biomarker amount $C_t = n$ uniformly distributed over a total volume $V_{tot}$ of plasma. If we sample from it a volume $V_s$, each biomarker unit is in the sample independent of the others with probability $p = \frac{V_s}{V_{tot}}$. Hence, the total amount of biomarker $X_t$ present in the sample is binomially distributed with parameters $n$ and $p$

$$\mathrm{P}(X_t = k \mid C_t = n) = \binom{n}{k} \left(\frac{V_s}{V_{tot}}\right)^k \left(1 - \frac{V_s}{V_{tot}}\right)^{n-k} \ . \tag{48}$$

The total amount of biomarker present in the plasma then follows by averaging over all the possible values of $C_t$

$$\mathrm{P}(X_t = k) = \sum_{n=k}^{\infty} \mathrm{P}(C_t = n)\, \mathrm{P}(X_t = k \mid C_t = n) \ . \tag{49}$$

The second term in the sum above is simply given by equation (48). As we noticed in the derivation of our asymptotic results, if $C_t$ follows a Poisson distribution, then due to the thinning property, $X_t$ follows a Poisson distribution as well. The expected value and variance of $X_t$ in terms of $C_t$ are given by

$$\mathrm{E}[X_t] = p\,\mathrm{E}[C_t]\,, \quad \mathrm{Var}(X_t) = p(1-p)\mathrm{E}[C_t] + p^2\,\mathrm{Var}(C_t)\ .$$

To condition on $A_t$ survival, we observe that

$$\mathrm{P}\left(X_t = k \mid \Omega_t\right) = \frac{\mathrm{P}(X_t = k) - \mathrm{P}(X_t = k, A_t = 0)}{1 - \delta_t}\ . \tag{50}$$

Of the two terms in the numerator, the first one coincides with equation (49), while the second one expands as

$$\begin{aligned} \mathrm{P}(X_t = k, A_t = 0) &= \sum_{n=k}^{\infty} \mathrm{P}(X_t = k, A_t = 0 \mid C_t = n)\,\mathrm{P}(C_t = n) \\ &= \sum_{n=k}^{\infty} \mathrm{P}(X_t = k \mid C_t = n)\,\mathrm{P}(A_t = 0 \mid C_t = n)\,\mathrm{P}(C_t = n)\ . \end{aligned}$$

The second equality follows from the fact that $X_t$ and $A_t$ are conditionally independent given $C_t$. Equation (50) thus becomes

$$\begin{aligned} \mathrm{P}(X_t = k \mid \Omega_t) &= \frac{\sum_{n=k}^{\infty} \mathrm{P}(X_t = k \mid C_t = n)\,\mathrm{P}(C_t = n)\,[1 - \mathrm{P}(A_t = 0 \mid C_t = n)]}{\mathrm{P}(\Omega_t)} \\ &= \frac{\sum_{n=k}^{\infty} \mathrm{P}(X_t = k \mid C_t = n)\,[\,\mathrm{P}(C_t = n) - \mathrm{P}(A_t = 0, C_t = n)]}{\mathrm{P}(\Omega_t)} \\ &= \frac{\sum_{n=k}^{\infty} \mathrm{P}(X_t = k \mid C_t = n)\,\mathrm{P}(C_t = n, \Omega_t)}{\mathrm{P}(\Omega_t)} \\ &= \sum_{n=k}^{\infty} \mathrm{P}(X_t = k \mid C_t = n)\,\mathrm{P}(C_t = n \mid \Omega_t)\ . \end{aligned}$$

#### References


1. R. Etzioni, *et al.*, *Nature Reviews Cancer* **3**, 243 (2003).
2. M. Song, B. Vogelstein, E. L. Giovannucci, W. C. Willett, C. Tomasetti, *Science* **361**, 1317 (2018).
3. S. Srivastava, *et al.*, *Nature Reviews Cancer* p. 1 (2019).
4. A. K. Mattox, *et al.*, *Science Translational Medicine* **11**, eaay1984 (2019).
5. R. L. Siegel, K. D. Miller, A. Jemal, *CA: A Cancer Journal for Clinicians* **70**, 7 (2020).
6. C. Bettegowda, *et al.*, *Science Translational Medicine* **6**, 224ra24 (2014).
7. A. M. Newman, *et al.*, *Nature Medicine* **20**, 548 (2014).
8. J. Phallen, *et al.*, *Sci Transl Med* **9** (2017).
9. J. D. Cohen, *et al.*, *Science* **357**, 926– (2018).
10. F. Mouliere, *et al.*, *Science Translational Medicine* **10** (2018).
11. S. Y. Shen, *et al.*, *Nature* **563**, 579 (2018).
12. S. Cristiano, *et al.*, *Nature* **570**, 385 (2019).
13. P. Ulz, *et al.*, *Nature Genetics* **48**, 1273 (2016).
14. J. J. Chabon, *et al.*, *Nature* (2020).
15. M. C. Liu, *et al.*, *Annals of Oncology* (2020).
16. J. C. Wan, *et al.*, *Nature Reviews Cancer* **17**, 223 (2017).
17. E. Heitzer, I. S. Haque, C. E. Roberts, M. R. Speicher, *Nature Reviews Genetics* **20**, 71 (2019).
18. H. Schwarzenbach, D. S. Hoon, K. Pantel, *Nature Reviews Cancer* **11**, 426 (2011).
19. A. M. Aravanis, M. Lee, R. D. Klausner, *Cell* **168**, 571 (2017).
20. S. S. Hori, S. S. Gambhir, *Science Translational Medicine* **3**, 109ra116 (2011).
21. S. S. Hori, A. M. Lutz, R. Paulmurugan, S. S. Gambhir, *Cancer Research* **77**, 2570 (2017).

22. B. Vogelstein, *et al.*, *Science* **339**, 1546 (2013).

23. M. Greaves, C. C. Maley, *Nature* **481**, 306 (2012).

24. D. W. Cescon, S. V. Bratman, S. M. Chan, L. L. Siu, *Nature Cancer* **1**, 276 (2020).

25. H. T. Winer-Muram, *et al.*, *Radiology* **223**, 798 (2002).

26. D. A. Rew, G. D. Wilson, *European journal of surgical oncology : the journal of the European Society of Surgical Oncology and the British Association of Surgical Oncology* **26**, 405 (2000).

27. C. Abbosh, *et al.*, *Nature* **545**, 446 (2017).

28. E. J. Moding, *et al.*, *Nature Cancer* **1**, 176 (2020).

29. A. M. Newman, *et al.*, *Nature Biotechnology* **34**, 547 (2016).

30. I. Bozic, *et al.*, *Proc Natl Acad Sci USA* **107**, 18545 (2010).

31. R. Durrett, *Branching process models of cancer* (Springer, 2015).

32. N. Beerenwinkel, R. F. Schwarz, M. Gerstung, F. Markowetz, *Systematic Biology* **64**, e1 (2015).

33. P. M. Altrock, L. L. Liu, F. Michor, *Nature Reviews Cancer* **15**, 730 (2015).

34. D. Wodarz, N. L. Komarova, *Dynamics of Cancer: Mathematical Foundations of Oncology* (World Scientific Publishing Co., Inc., Singapore, 2014).

35. B. R. McDonald, *et al.*, *Science Translational Medicine* **11**, eaax7392 (2019).

36. J. Tie, *et al.*, *Science Translational Medicine* **8**, 346ra92 (2016).

37. K. H. Khan, *et al.*, *Cancer Discovery* **8**, 1270 (2018).

38. T. Reinert, *et al.*, *JAMA Oncology* (2019).

39. A. A. Chaudhuri, *et al.*, *Cancer Discovery* **7**, 1394 (2017).

40. J. C. M. Wan, *et al.*, *bioRxiv* p. 759399 (2019).

41. NCI, Surveillance, epidemiology, and end results (seer) research data 1973-2015 - ascii text data (2018).

42. I. Martincorena, P. J. Campbell, *Science* **349**, 1483 (2015).

43. A. Yokoyama, *et al.*, *Nature* **565**, 312 (2019).

44. A. P. Makohon-Moore, *et al.*, *Nature* **561**, 201 (2018).

45. P. Razavi, *et al.*, *Nature Medicine* **25**, 1928 (2019).

46. A. McWilliams, *et al.*, *New England Journal of Medicine* **369**, 910 (2013).

47. The National Lung Screening Trial Research Team, *New England Journal of Medicine* **368**, 1980 (2013).

48. F. C. Detterbeck, D. J. Boffa, A. W. Kim, L. T. Tanoue, *Chest* **151**, 193 (2017).

49. C. A. Clarke, *et al.*, *Cancer Epidemiology and Prevention Biomarkers* (2020).

50. N. L. S. T. R. Team, *New England Journal of Medicine* **365**, 395 (2011). PMID: 21714641.

51. H. J. de Koning, *et al.*, *New England Journal of Medicine* (2020).

52. A. P. Makohon-Moore, C. A. Iacobuzio-Donahue, *Nature Reviews Cancer* **16**, 553 (2016).

53. B. M. Lang, J. Kuipers, B. Misselwitz, N. Beerenwinkel, *PLOS Computational Biology* **16**, e1007552 (2020).

54. M. E. Hochberg, F. Thomas, E. Assenat, U. Hibner, *Evolutionary Applications* **6**, 134 (2013).

55. H. J. de Koning, *et al.*, *Annals of Internal Medicine* **160**, 311 (2014).

56. K. Curtius, A. Dewanji, W. D. Hazelton, J. H. Rubenstein, E. G. Luebeck, *bioRxiv* (2020).

57. M. D. Ryser, *et al.*, *Journal of the National Cancer Institute* **108**, djv372 (2016).

58. K. Lahouel, *et al.*, *Proc Natl Acad Sci USA* **117**, 857 (2020).

59. K. B. Athreya, P. Ney, *Branching Processes* (Dover Publications, 2004).

60. M. Kimmel, D. Axelrod, *Branching Processes in Biology* (Springer, New York, 2015).

61. V. H. Teixeira, *et al.*, *Nature Medicine* **25**, 517 (2019).

62. I. Martincorena, *et al.*, *Science* **362**, 911 (2018).

63. L. Allen, *An introduction to stochastic processes with applications to biology* (Boca Raton, FL: Chapman & Hall/CRC, 2011), second edn.

64. V. G. Kulkarni, *Modeling and analysis of stochastic systems* (Chapman and Hall/CRC, 2016).

65. F. W. Steutel, K. Van Harn, *Infinite divisibility of probability distributions on the real line* (CRC Press, 2003).

66. D. Cheek, T. Antal, *et al.*, *The Annals of Applied Probability* **28**, 3922 (2018).

67. D. V.-J. D.J. Daley, *Introduction to the Theory of Point Processes* (Springer New York, 2006).

68. T. Antal, P. L. Krapivsky, *J. Stat. Mech. Theory Exp.* **2010**, P07028, 22 (2010).

69. T. Antal, P. L. Krapivsky, *Journal of Statistical Mechanics P08018* (2011).

70. A. Ronveaux, *Heun's Differential Equations* (Oxford University Press, 1995).

71. O. V. Motygin, *2018 Days on Diffraction (DD)* (IEEE, 2018).

72. M. Abramowitz, I. Stegun, *Handbook of Mathematical Functions: With Formulas, Graphs, and Mathematical Tables*, Applied mathematics series (Dover Publications, 1964).

73. J. Abate, W. Whitt, *Operations Research Letters* **12**, 245 (1992).

74. G. L. Choudhury, D. M. Lucantoni, W. Whitt, *The Annals of Applied Probability* **4**, 719 (1994).
